# Non-coding *NFKBIZ* 3′ UTR mutations promote cell growth and resistance to targeted therapeutics in diffuse large B-cell lymphoma

**DOI:** 10.1101/2021.05.22.445261

**Authors:** Sarah E. Arthur, Jie Gao, Shannon Healy, Christopher K. Rushton, Nicole Thomas, Laura K. Hilton, Kostiantyn Dreval, Jeffrey Tang, Miguel Alcaide, Razvan Cojocaru, Anja Mottok, Adèle Telenius, Peter Unrau, Wyndham H. Wilson, Louis M. Staudt, David W. Scott, Daniel J Hodson, Christian Steidl, Ryan D. Morin

## Abstract

Amplifications and non-coding 3′ UTR mutations affecting *NFKBIZ* have been identified as recurrent genetic events in diffuse large B-cell lymphoma (DLBCL). We confirm the prevalence and pattern of *NFKBIZ* 3′ UTR mutations in independent cohorts and determine they are enriched in the ABC subtype as well as the recently described novel BN2/C1/NOTCH2 classes of DLBCL. Presently, the effects of and mechanism by which non-coding mutations can act as cancer drivers has been relatively unexplored. Here, we provide a functional characterization of these non-coding *NFKBIZ* 3′ UTR mutations. We demonstrate that the resulting elevated expression of IκB-ζ confers growth advantage in DLBCL cell lines and primary germinal center B-cells as well as nominate novel IκB-ζ target genes with potential therapeutic implications. The limited responses to targeted treatments in DLBCL, particularly those targeting the NF-κB axis, led us to investigate and confirm that *NFKBIZ* 3′ UTR mutations affect response to therapeutics and suggest it may be a useful predictive biomarker.

**Statement of Significance:** Through functional characterization we reveal that non-coding *NFKBIZ* 3′ UTR mutations are a common driver in DLBCL, and mutation status may be a relevant biomarker to predict poor response to therapeutics targeting the NF-κB pathway.

## Introduction

Diffuse large B-cell lymphoma (DLBCL) represents the most common form of non-Hodgkin lymphoma worldwide. Historically, DLBCL has been divided into two molecular subgroups through gene expression profiling (GEP), namely the activated B-cell-like (ABC) and germinal centre B-cell-like (GCB) subgroups^1^. These distinct cell-of-origin (COO) subgroups are generally appreciated as arising through unique mechanisms and rely on activation of specific oncogenic pathways. ABC DLBCLs are characterized by constitutive activation of the nuclear factor-κB (NF-κB) signaling pathway, which can often be explained by mutations in genes such as *MYD88, CARD11, CD79A/B* or *TNFAIP3*^2–4^. Patients with ABC DLBCL generally experience an inferior clinical response to standard-of-care treatment. Although many alterations leading to NF-κB pathway activation have been described, there are a substantial number of ABC cases with no clear genetic explanation for their NF-κB activation.

Recent work has shifted attention to the use of genetic features for classifying DLBCLs with shared biology that may inform on therapeutic vulnerabilities^5–7^. Despite the newly described subtypes representing a more granular division of this disease, there remain discrepancies between classification systems and many tumors still remain unclassifiable. This suggests that further refinement is needed to classify and predict treatment options for all DLBCL patients^8^. To enable the identification and validation of new treatment options and stratify DLBCL patients accordingly in relevant clinical trials, it is important to fully understand the molecular underpinnings of this heterogenous disease. Thus, the discovery and characterization of mutations which could act as relevant biomarkers for targeted treatments is ongoing. This includes the characterization of non-coding regulatory mutations that can represent *bona fide* driver mutations by dysregulating gene expression.

Our group recently described a pattern of non-coding somatic mutations affecting the 3′ untranslated region (UTR) of the *NFKBIZ* gene^9^, which has since been confirmed by other studies^10^. *NFKBIZ* encodes for the IκB-ζ protein, an atypical member of the nuclear IκB protein family involved in the NF-κB signaling pathway^11^. Expression of IκB-ζ is barely detectable in resting cells, but rapidly induced upon stimulation of Toll-like receptors^11^ and other stimuli feeding into the NF-κB pathway (e.g. LPS and interleukins^12^). IκB-ζ has been shown to regulate NF-κB signaling, however it has been implicated in both repression^11^ and, more recently, activation^12^ of this pathway. Recent studies suggest that it may inhibit and activate distinct subsets of NF-κB target genes^13^. Specifically, IκB-ζ may confer transactivating potential to p50 and p52 homodimers to promote transcription^14^, but may also block the formation of p65 heterodimers, thus inhibiting their function^15^. *Nogai et al*. found that IκB-ζ is highly expressed in most ABC DLBCL cell lines and it was previously discovered that the *NFKBIZ* locus is amplified in 10% of ABC DLBCLs^16^.

Given the high expression rates of *NFKBIZ* in ABC DLBCLs, we hypothesized that *NFKBIZ* 3′ UTR mutations act to promote IκB-ζ expression by stabilizing *NFKBIZ* mRNA. Consistent with this notion, DLBCL cell lines with naturally occurring *NFKBIZ* 3′ UTR mutations expressed elevated *NFKBIZ* mRNA and IκB-ζ protein levels relative to other DLBCL lines^9^. Here, we provide a functional characterization of the mechanism and effect of non-coding *NFKBIZ* 3′ UTR mutations in DLBCL and suggest their potential utility as a predictive biomarker for response to targeted therapeutics.

## Results

### *NFKBIZ* 3′ UTR mutations and amplifications are enriched in ABC DLBCL and the BN2 and A53 subtypes

Since the first description of non-coding mutations in the *NFKBIZ* 3′ UTR in DLBCL^9^, multiple studies have released sequencing data from additional cohorts but focused on protein coding region alterations^5,6^. We reanalyzed these datasets to search for non-coding *NFKBIZ* 3′ UTR mutations and compared their general prevalence among cohorts and within each of the molecular and genetic subgroups. A comparison of the mutations from our previously published dataset (Arthur *et al*.)^9^ with new mutation calls from Schmitz *et al*.^6^ and Chapuy *et al*.^5^ revealed a similar pattern of *NFKBIZ* mutations, with small deletions and single-nucleotide variants (SNVs) being the most common mutations affecting the 3′ UTR (**Figure 1A, D** and **E**). When amplifications and 3′ UTR mutations are considered, *NFKBIZ* was among the top seven recurrently altered genes in DLBCL (**Figure 1A**). Although coverage varied, the majority of patients in the Schmitz cohort had sufficient sequencing depth for the most commonly mutated region of the 3′ UTR to be covered. The Chapuy dataset had consistently low coverage of the 3′ UTR, which limited our ability to detect mutations in those tumors (**Supplemental Figure S1**). Interestingly, *NFKBIZ* 3′ UTR mutations (102/1006 patients) and *MYD88*^L265P^ hotspot mutations (134/1006 patients) were never observed in the same tumor (**Figure 1B**). We found that all mutations in the *MYD88* and *NFKBIZ* genes were significantly mutually exclusive (P < 1.25⨯10^-3^) within ABC DLBCLs, where these mutations are predominantly found (**Figure 1C**). Overall, *NFKBIZ* is mutated (UTR or amplification) in 17% of DLBCLs and 24% of the ABC subtype (**Figure 1D, Supplemental Table S1**), however this is likely an underestimate due to the variability in coverage between datasets (**Figure 1E, Supplemental Table S1**). We also searched for *NFKBIZ* 3′ UTR mutations in numerous other cohorts with whole genome sequencing (WGS) data and found mutations in DLBCL^9,17,18^, follicular lymphoma (FL)^18,19^, and from published and unpublished Burkitt lymphoma (BL) cohorts from the Burkitt Lymphoma Genome Sequencing Project (BLGSP)^20^ (**Supplemental Table S2 and S3**). This confirmed our previous observation that these mutations are infrequently observed in other B-cell neoplasms. The mutations in DLBCL and the other malignancies show a similar pattern to that observed among exome-based data sets that comprised the core of our analysis.

**Figure 1.**
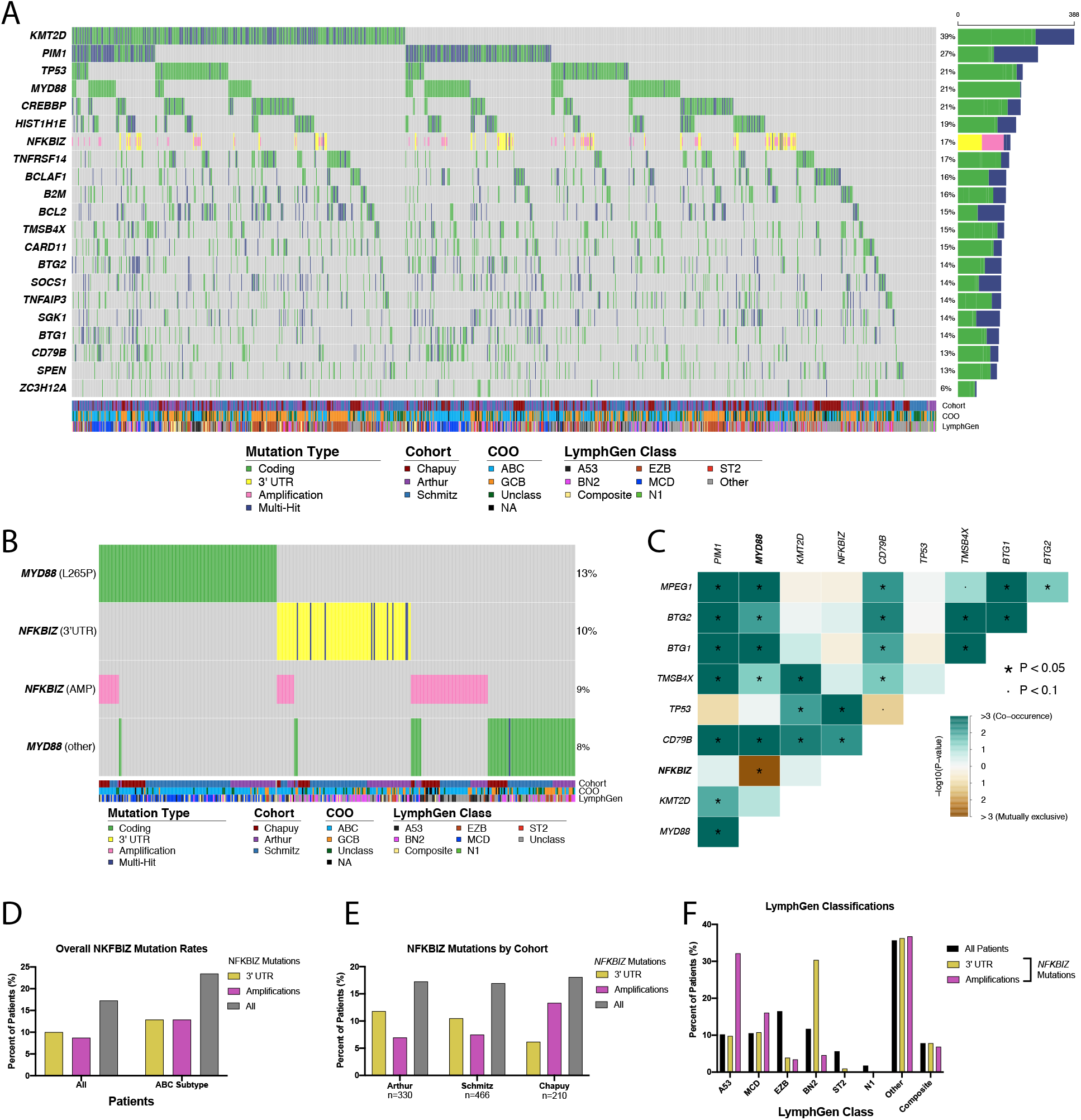
*NFKBIZ* mutation prevalence and mutual exclusivity in DLBCL. (**A**) Oncoplot showing the top 20 most commonly mutated genes within the Arthur^9^ and Chapuy^5^ and Schmitz^6^ cohorts. The gene *ZC3H12A* was added to show its prevalence compared to the top genes. Cohort, cell of origin (COO) and LymphGen classifications are shown in tracks at the bottom. (**B**) Oncostrip showing *MYD88* and *NFKBIZ* mutations split up by mutation type. *MYD88* hotspot (L265P) mutations never co-occur with *NFKBIZ* 3′ UTR mutations. (**C**) Somatic interactions plot for ABC DLBCL cases showing the top 10 genes with mutations that co-occur (green) and are mutually exclusive (brown) (*P<0.05, •P<0.1, pair-wise Fisher’s Exact test). *NFKBIZ* and *MYD88* mutations are significantly mutually exclusive. (**D**) Prevalence of all *NFKBIZ* mutations including 3′ UTR and amplifications in all DLBCLs and only considering ABC DLBCLs. (**E**) Prevalence of *NFKBIZ* mutations within each of the analyzed cohorts. (**F**) Prevalence of *NFKBIZ* mutations within each of the LymphGen classes compared to the breakdown of all cases within each subtype.

We next explored the relationship between *NFKBIZ* mutations and the newly described genetic subgroups of DLBCL^5–7^. Notably, gains/amplifications of 3q (location of *NFKBIZ*) and/or focal amplifications of *NFKBIZ* were genetic features used in some of these classifiers and are typically associated with cases assigned to the MCD subgroup^6^ (also corresponding to the C5^5^ and MYD88^7^ groups) but more recently, with the addition of extra subgroups to LymphGen, they are also enriched in the newly described A53 subgroup, which is mostly defined by copy number features^21^. In contrast, *NFKBIZ* 3′ UTR mutations were not considered when defining these classification systems. Using LymphGen^22^ to assign samples from all three cohorts to genetic subgroups, we observed a similar pattern of classification (**Supplemental Table S4**), with a high proportion of *NFKBIZ* amplification cases classified as A53. Interestingly, *NFKBIZ* 3′ UTR mutations were most commonly observed in BN2 cases (**Supplemental Table S3 and S4**). BN2 is characterized by mutations in other components of the BCR-dependent NF-κB pathway (*PRKCB, BCL10, TNFAIP3, TNIP1*). This observation is consistent with the notion that *NFKBIZ* 3′ UTR mutations could be functionally distinct from amplifications and provide a selective advantage specifically in BN2 tumors. The mutual exclusivity of mutations in *MYD88* and *NFKBIZ* may also explain the paucity of MCD tumors with these mutations. This highlights that distinct genetic alterations, each contributing to deregulation of NF-κB signaling, are appropriately segregated by current classification methods but the presence of some NFKBIZ-mutant tumours in the “unclassified” group should be noted.

### *NFKBIZ* 3′ UTR Secondary Structure

Post-transcriptional regulation of genes can involve interactions between conserved sequences or secondary structures and either miRNAs or RNA-binding proteins (RBPs), which can lead to either stabilization or destabilization of transcripts. We sought to clarify the structural features of the *NFKBIZ* 3′ UTR that are commonly mutated. Using a combination of RNA secondary structure prediction tools^23–26^ (**Supplemental Figure S2**) accompanied by RNA probing assays (**Supplemental Figure 2A-C**), we constructed a refined model of the *NFKBIZ* 3′ UTR secondary structure (**Figure 2D-E**). To do this, predicted structures were compared with the results from an RNA-probing assay which determined guanine (G) bases that were paired (protected) or unpaired (exposed) under physiological conditions. Through this comparison we selected the most comprehensive model that included structural features predicted by multiple tools and fitted with experimental data. The structure comprises two novel stem-loops (SL1 and SL2) and three previously described stem-loops (SL3, SL4, and SL5)^27–30^.

**Figure 2.**
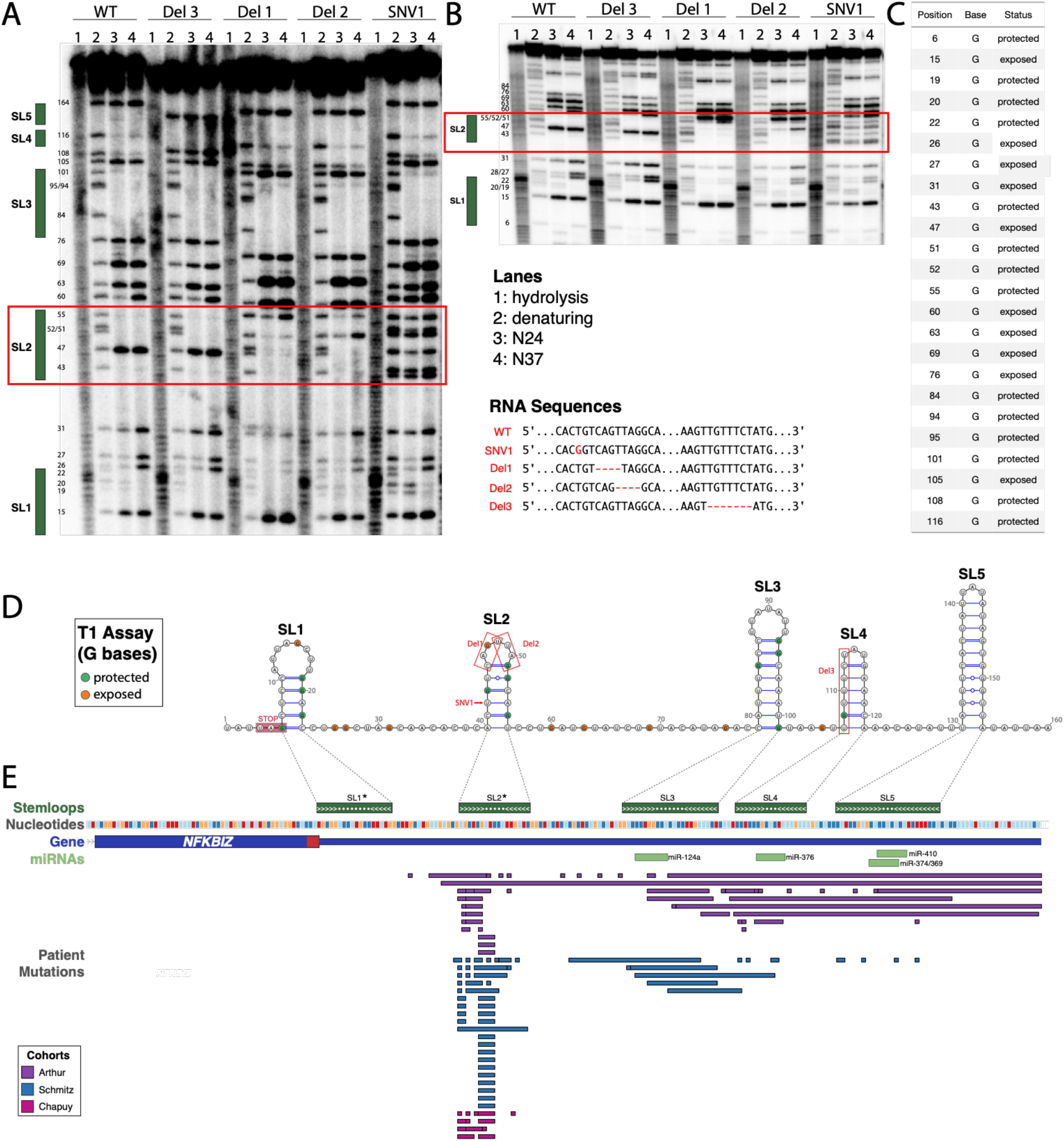
Novel stem-loop structure predicted in the *NFKBIZ* 3′ UTR. (**A**) T1 RNase probing assays (single-stranded guanine bases) were performed with four variants of the *NFKBIZ* 3′ UTR RNA (WT, SNV1, Del1, Del2 and Del3). Four conditions were tested: Hydrolysis (all bases cut), denaturing (all G’s cut), N24 (native conformation at 24C) and N37 (native conformation at 37 ^o^C). Gels were run for different amounts of time to resolve both (**A**) 3′ and (**B**) 5′ ends of the RNA sequence. Green boxes beside each gel denote the location of suspected stem-loop structures (SL1-SL5). The red box shows the region containing SL2 and highlights the differences in structure between WT and mutants at this position. (**C**) Summary table of which G bases were protected (not cut – part of a structure) or exposed (cut – not part of a structure). This was used as a basis to confirm predicted structure of the region in panel D. (**D**) Consensus predicted structure of the *NFKBIZ* 3′ UTR. The final prediction is a representation of the structure with the most overlap between RNA folding prediction algorithms and that is still consistent with experimental T1 assay results. Protected and exposed guanine (G) bases from the T1 assay are shown in green and orange. Location of mutations in RNA used for probing assays are shown in red (SNV1, Del1, Del2 and Del 3). (**B**) 3′ UTR region of *NFKBIZ* showing stem-loop structures in dark green (novel SLs denoted by *), dashed lines connect SLs to the 2D structure from panel A, nucleotides in this region (dark blue: T, light blue: A, red: G, orange: C), the gene body is shown in dark blue with the last exon, stop codon (red) and 3′ UTR (untranslated region), the location of previously predicted microRNAs (miRNAs) are shown in light green and the location of patient mutations from three cohorts in Figure 1A (purple: Arthur, blue: Schmitz and pink: Chapuy).

This region of the *NFKBIZ* 3′ UTR contains four known miRNA binding sites (**Figure 2E**), all found in the three known distal stem-loops (SL3, SL4 and SL5)^31,32^. Each of SL3, SL4 and SL5 have also been described as specific targets of the RBPs Regnase-1 and Roquin^27–30,33^. These RBPs are important for the degradation and translational silencing of many inflammation-related mRNAs such as the pro-inflammatory cytokines TNFα and IL-6. The stem-loop SL2, the region most densely affected by mutations, is similar in structure to other targets of Regnase-1 and Roquin^34–36^ and therefore we postulate that it may also be targeted by these RBPs. The distinct lack of mutations affecting SL1 is also notable and may imply a distinct function for this region. The enrichment of mutations affecting SL2 suggests that perturbations to this structure may have the most drastic effect on *NFKBIZ* (and IκB-ζ) expression levels. Through RNA-probing assays, we noted that the *in vitro* cleavage patterns of RNA representing four separate patient-derived mutations (Del1, Del2, Del3 and SNV1) was each distinct from WT RNA. We confirmed that the three mutations affecting the region containing SL2 (Del1, Del2 and SNV1) disrupted the structure of SL2 (**Supplemental Figure 2A-B**). The extent to which mutations disrupt RBPs targets and/or miRNA binding sites is likely variable and dependent on the exact location of each mutation. Consistent with this model wherein *NFKBIZ* deregulation is achieved by modulating interactions with RBPs, we note the presence of recurrent non-silent mutations affecting the gene encoding the RBP Regnase-1 (*ZC3H12A*) in DLBCL (**Figure 1A**).

### CRISPR-induced *NFKBIZ* 3′ UTR mutations lead to elevated mRNA and protein levels

Using the CRISPR-Cas9 system, we introduced mutations into the *NFKBIZ* 3′ UTR of two DLBCL cell lines, one ABC and one GCB (U2932 and WSU-DLCL2, respectively). As the majority of patient mutations affect SL2, we designed a guide targeting this region. To aid in comparing their potential role as driver mutations, we also designed a guide to target one of the other stem-loops (SL3) that is mutated at a lower incidence. Eight clones with different SL2 mutations were obtained from the WSU-DLCL2 line (**Figure 3A-C**) including one 1 bp insertion, six small deletions and one large deletion. Five clones with mutations targeting SL2 and SL3 were obtained in the U2932 cell line (Figure **3A-B,D**). WSU-DLCL2 is hemizygous at this locus and therefore mutants had no WT allele, whereas some U2932 clones had mutations on both alleles and the remainder had one mutant and one WT allele.

**Figure 3.**
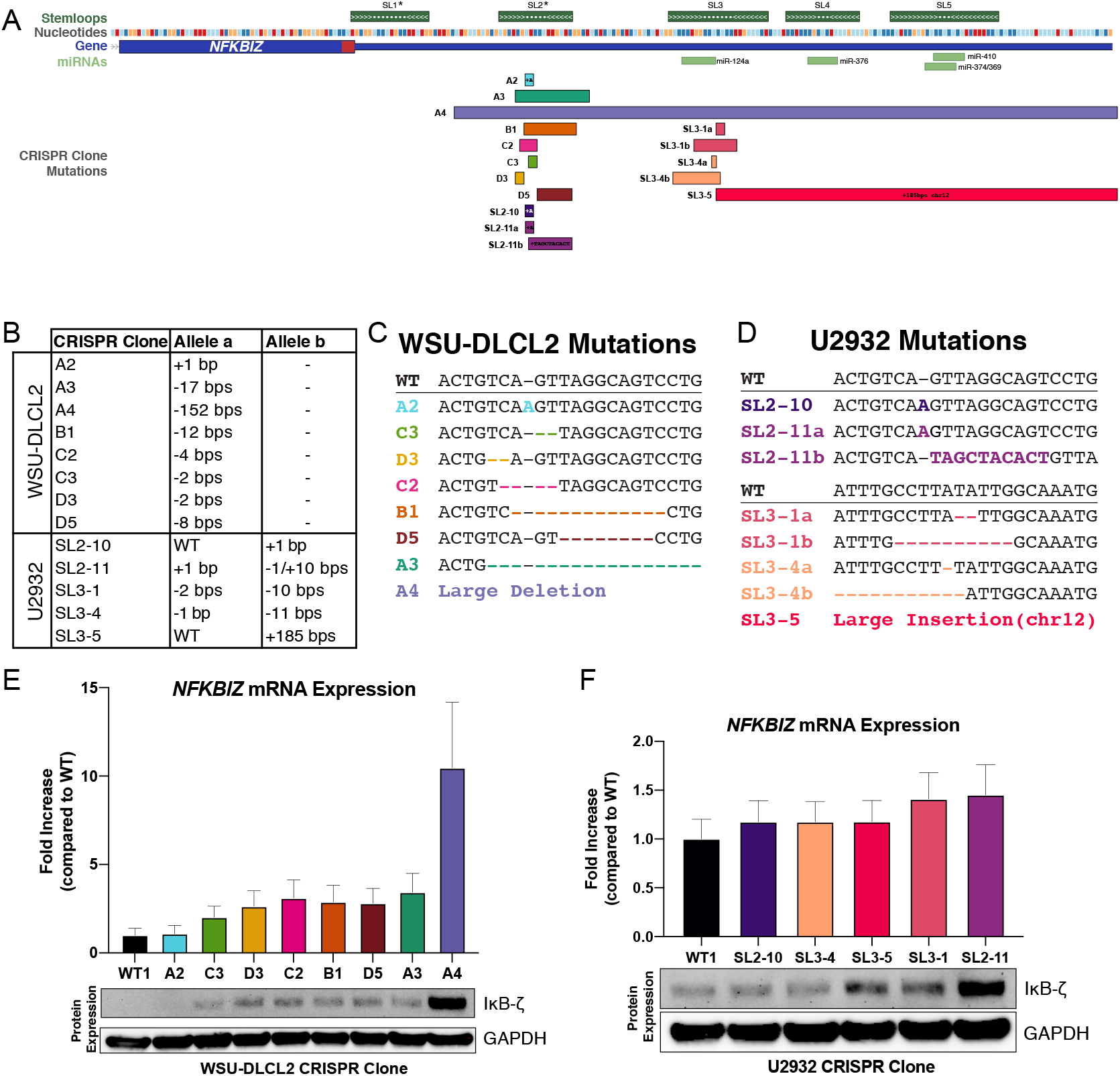
CRISPR-induced *NFKBIZ* 3′ UTR mutations in DLBCL cell lines. (**A**) Location of mutations seen in CRISPR clones from two cell lines (WSU-DLCL2 and U2932) relative to SL structures described in Figure 2. (**B**) Table showing exact mutation sizes in each allele where relevant. (**C**) Sequence of mutations seen in WSU-DLCL2 clones targeting stem-loop 2 region. (**D**) Sequence of mutations seen in U2932 clones targeting stem-loop 2 and stem-loop 3 regions. *NFKBIZ* mRNA (top) and IκB-ζ protein expression (bottom) of (**E**) WSU-DLCL2 and (**F**) U2932 CRISPR clones compared to WT from droplet digital PCR (ddPCR) and western blot, respectively.

We used droplet digital PCR (ddPCR) and western blot analysis respectively to determine *NFKBIZ* mRNA and IκB-ζ protein expression in WT and mutant clones (**Figure 3E-F**). Most lines with mutations expected to disrupt structural elements in the UTR exhibited elevated mRNA that corresponded to elevated protein levels in both cell lines. In the WSU-DLCL2 clones, there was no detectable IκB-ζ protein in the WT clone by western blot, but all deletion clones had detectable bands corresponding to IκB-ζ protein. Although the WT U2932 clone had detectable IκB-ζ protein, some of the mutant clones still showed increases in band intensity for IκB-ζ protein. One exception in both lines were clones representing the same 1 bp insertion affecting a predicted loop (WSU-DLCL2 A2 and U2932 SL2-10), which was not predicted to disrupt the SL structure. This insertion was also never observed in patient samples and thus is unlikely to act as a driver mutation. Consistent with this interpretation, neither of these clones had noticeably elevated mRNA or IκB-ζ protein levels.

The clone with the most striking change in *NFKBIZ* expression was WSU-DLCL2 A4, which harbors a deletion that removes four of five stem-loops (SL2-SL5). We suspect that deletion of this entire region disrupts multiple regulatory features, leading to a more drastic increase of expression. Although we have seen structural variations comprising large deletions of the *NFKBIZ* 3′ UTR, small indels (especially deletions) in SL2 are far more frequent in patients. This could be explained by mutational processes that favour the introduction of small indels or may imply that the higher increase of expression accomplished by large deletions is not necessary or has additional consequences causing a reduced selective advantage.

### IκB-ζ expression confers a selective growth advantage

To investigate the effect of elevated IκB-ζ expression caused by *NFKBIZ* 3′ UTR mutations on cell growth, we performed competitive growth assays comparing the parental (WT) cell line to mutant clones both *in vitro* (**Figure 4A**) and *in vivo* (**Figure 4D**). WT and mutant WSU-DLCL2 lines were pooled and grown for 8-9 passages in cell culture or injected into a mouse and left to grow into a tumor xenograft. Based on the relative proportion of each mutation in the population, all deletion-bearing clones outgrew WT (**Figure 4B**), with the A4 clone showing the highest representation. This clone exhibited the highest IκB-ζ expression levels, consistent with IκB-ζ expression conferring a selective growth advantage in culture as a function of expression level. In a separate experiment we compared WT individually to a subset of mutants and confirmed that each mutant clone can individually out-compete WT *in vitro* (**Figure 4C**). This highlights that the increased expression levels of IκB-ζ from small deletions are sufficient to cause a growth advantage, and large deletions (similar to A4), are not necessary to observe this effect. A similar growth advantage was observed *in vivo* through xenograft experiments. In mice, the A4 clone represented the majority of cells constituting the tumor at the endpoint (**Figure 4F**). Interestingly, when separately injected into mouse models, WT and A4 tumors grew at a similar rate, however it appeared that the A4 group may have seeded detectable tumors slightly earlier (**Supplemental Figure S3**).

**Figure 4.**
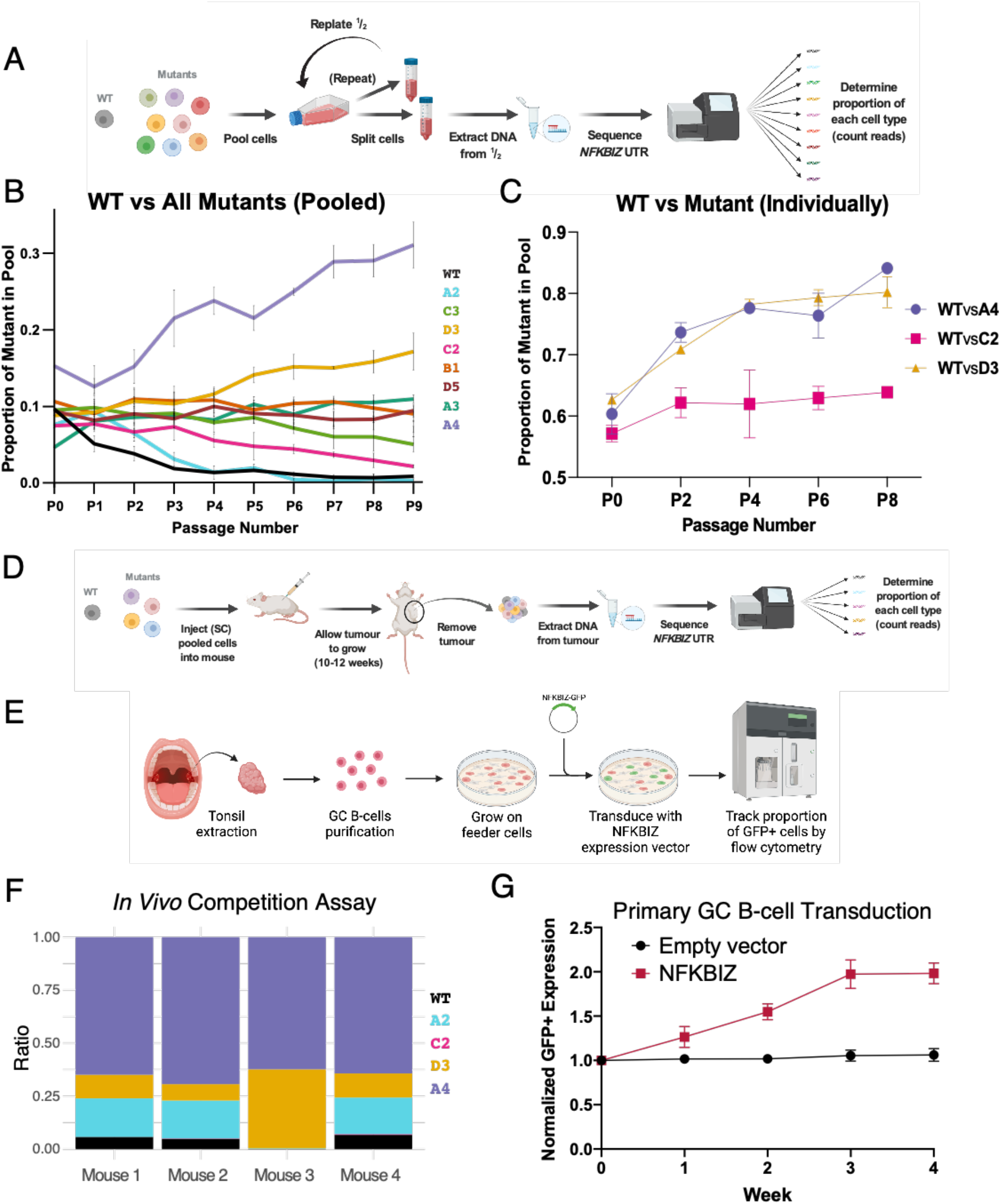
*NFKBIZ* expression provides a selective growth advantage to cells. (**A**) Schematic of *in vitro* competitive growth experiments. (**B**) Proportion of WT and mutant clones in the competitive growth assay pool of all clones grown together over multiple passages. DNA was extracted from cell pellets at each time point and sequenced to determine proportion of the pool made up by each clone. Each line represents the growth of a clone within one pooled experiment with three replicates. (**C**) Proportion of mutant clones when grown separately against WT in culture over multiple passages. DNA was extracted at each time point and ddPCR was used to differentiate WT and mutant sequences to infer proportion of the pool. Each line represents an individual experiment of three replicates. (**D**) Schematic of *in vivo* competitive growth experiments. (**E**) Schematic of primary germinal center B-cell *NFKBIZ* expression vector transduction experiments. (**F**) Proportion of WT or mutant cells in xenograft tumor after grown to endpoint. DNA was extracted from the tumor at endpoint and sequenced to determine the proportion of each clone represented in the tumor. (**G**) Primary human germinal center B cells were purified from discarded tonsil tissue, cultured on irradiated YK6-CD40Lg-IL21 feeders and immortalized with BCL6-t2A-BCL2. Cells were then transduced with either NFKBIZ-IRES-GFP or control empty vector-GFP. The frequency of GFP-positive cells was measured by flow cytometry 4 days after transduction (week 0) and at weekly intervals thereafter. The frequency of GFP-positive cells, normalized to week 0, is shown for NFKBIZ and control cultures over a 4-week time course. Points represent mean and standard error of nine replicate cultures from six separate donors.

As cancer cell lines individually harbor advantageous mutations, we next sought to evaluate the effects of IκB-ζ expression in a genetically normal background. We directly determined the effect of increased IκB-ζ expression in primary germinal centre B-cells through ectopic expression. Primary human germinal centre B-cells immortalized with BCL6-t2A-BCL2 were transduced with a *NFKBIZ* overexpression vector or a control empty vector (**Figure 4E**). The growth of transduced cells was determined by measuring GFP-positive cells over time (**Figure 4G** and **Supplemental Figure S4**). Compared to cells transduced with a control vector, *NFKBIZ* overexpressing cells exhibited a significant competitive growth advantage *in vitro*.

### Candidate IκB-ζ target genes and pathways affected by *NFKBIZ* expression

We hypothesized that the selective growth advantage of cells with elevated IκB-ζ expression observed *in vitro* and *in vivo* is afforded by the activation of target genes and pathways regulated by IκB-ζ. To elucidate the genes and pathways with expression changes associated with IκB-ζ overexpression, we performed RNA-seq on a subset of *NFKBIZ* CRISPR-mutant clones and WT replicates. The WSU-DLCL2 parental line is GCB based on the COO classification and is classified as EZB by LymphGen. By comparing genes differentially expressed between WSU-DLCL2 WT and mutant clones, we identified 34 genes suspected to be induced or suppressed by IκB-ζ (**Figure 5A**). These include potential novel IκB-ζ targets *HCK, CD274* (PD-L1) as well as genes known to be affected by *MYD88* knockdown (*ARG2, LGALS3, CCR7, HTR3A*)^37^. Next, using gene set enrichment analysis (GSEA), we identified multiple hallmark pathways with significant changes in expression (**Figure 5B**). As expected, this included pathways relating to NF-κB signaling. Analysis of the gene expression changes in *NFKBIZ* 3′ UTR mutant lines revealed that a GCB line (WSU-DLCL2), following introduction of *NFKBIZ* 3′ UTR mutations, exhibited higher expression of pathways generally associated with ABC DLBCL such as TNF signaling via NF-κB and lower expression of GCB associated pathways such as MYC and MTOR (**Figure 5B-D**). This is consistent with *NFKBIZ* overexpression contributing, in part, to the ABC gene expression program.

**Figure 5.**
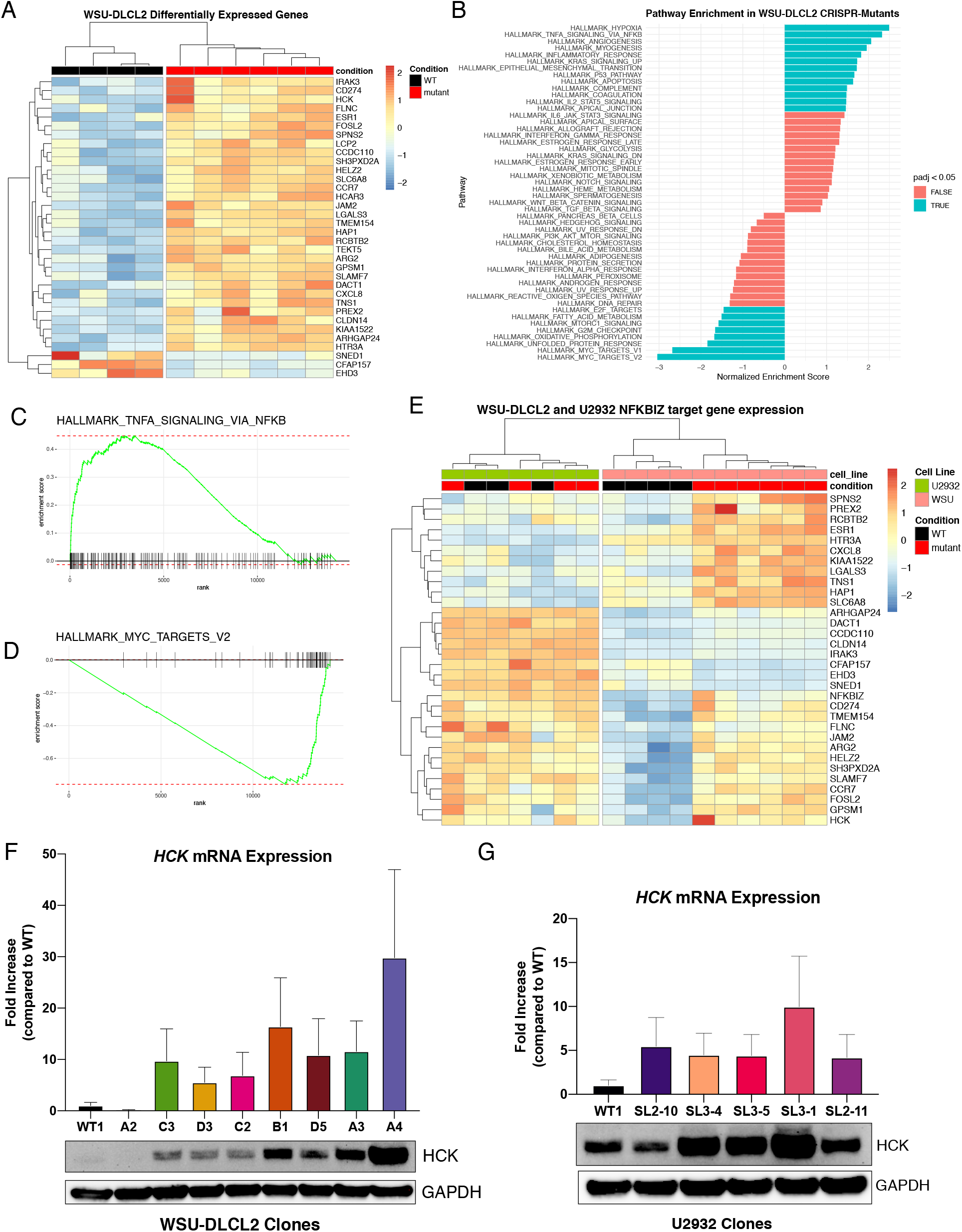
RNA-sequencing of CRISPR clones revealed novel targets and pathways activated by IκB-ζ. (**A**) Top genes differentially expressed between WSU-DLCL2 WT and *NFKBIZ* CRISPR-mutant clones. (**B**) Top pathways enriched in WSU-DLCL2 CRISPR mutant clones. Gene set signatures of top pathways upregulated (**C**) TNFα signaling via NF-κB and downregulated (**D**) MYC hallmark targets V2. (**E**) Expression of differentially expressed genes discovered in WSU-DLCL2 CRISPR clone analyses shown in both sets of CRISPR clones: WSU-DLCL2 and U2932. No differentially expressed genes were found between U2932 WT and CRISPR-mutant clones. HCK mRNA (top) and protein expression (bottom) of (**F**) WSU-DLCL2 and (**G**) U2932 CRISPR clones compared to WT from droplet digital PCR (ddPCR) and western blot, respectively.

In contrast to WSU-DLCL2, U2932 has been classified as ABC (unclassified by LymphGen) and has high basal expression of many NF-κB targets and genes considered to be ABC-like but lacks mutations typical among ABC/MCD DLBCL such as *MYD88, CD79B* or *NFKBIZ* (**Supplemental Figure S5**). Despite the upregulation of IκB-ζ in U2932, we did not identify any significantly differentially expressed genes between WT and the mutant clones. We also directly assessed the effect of *NFKBIZ* 3′ UTR mutations in U2932 cells line by comparing the expression of candidate IκB-ζ targets determined through analyses of the WSU-DLCL2 RNA-seq data between the U2932 WT and mutant clones (**Figure 5E**). *NFKBIZ* itself and, as expected, many of these candidate target genes were highly expressed in all U2932 lines including the parental line. In aggregate, we interpret these results as the lack of further increase in gene expression in a cell line that already has high activity of NF-κB signaling.

We next sought to validate the induction of *HCK* as a novel IκB-ζ target gene. We confirmed that the increase of *HCK* seen in RNA-sequencing was consistent with increased HCK mRNA through ddPCR and protein expression through western blot in CRISPR-mutant cell lines (both WSU-DLCL2 and U2932 clones) (**Figure 5F** and **5G**). Increased HCK expression was generally proportional to the level of IκB-ζ expression, with mutant clones showing a similar increase of both IκB-ζ and HCK protein levels. Given the role of HCK in other lymphoid cancers, it is appealing to consider whether the IκB-ζ induction of HCK may be contributing to the phenotype of *NFKBIZ*-mutant cells.

### *NFKBIZ* 3′ UTR mutations impart differential vulnerability to targeted therapeutics

Various targeted agents that have been or are currently being evaluated for their efficacy in treating ABC DLBCL have had limited success in clinical trials, with most providing therapeutic benefit in only a minority of patients^38–46^. Considering the genetic heterogeneity of DLBCL, it is important to understand the genetics underlying each tumor such that we may identify appropriate therapeutics that specifically target oncogenic pathways operative in that cancer. To this end, we used our engineered cell lines to investigate the effect of *NFKBIZ* overexpression on the efficacy of four targeted therapeutics that are under evaluation for application to DLBCLs, particularly those that either directly or indirectly perturb NF-κB signaling.

Specifically, we explored the *in vitro* toxicity (IC50) of ibrutinib (a BTK inhibitor)^42,45^, idelalisib (a PI3Kδ inhibitor)^47,48^, masitinib (a pan-SRC kinase inhibitor)^49^ and bortezomib (a proteasome inhibitor)^44,50^. Because the IκB-ζ protein acts downstream of many of these drug targets in the NF-κB pathway, we hypothesized that these mutations may confer resistance to some of these agents. Consistent with this hypothesis we found that the WSU-DLCL2 CRISPR-mutant cell lines had a significantly higher IC50 than the WT line for each of ibrutinib, idelalisib and masitinib. Interestingly, there was no significant difference in IC50 between WT and mutant lines treated with bortezomib (**Figure 6A-D**). The three drugs with differential effects on mutant lines all target proteins upstream of IκB-ζ in the NF-κB pathway (**Figure 7**), which could explain why an activating mutation in the *NFKBIZ* gene would afford cells resistance to these drugs. Bortezomib acts to inhibit degradation of the IκB proteins that inhibit NF-κB proteins from translocating into the nucleus where they form complexes with IκB-ζ. By blocking the release of NF-κB proteins, IκB-ζ is not able to confer trans-activating potential and bortezomib may be effective even in *NFKBIZ* mutant cases. The U2932 parental cell line is already known to be resistant to many of these drugs^51,52^ and therefore was not evaluated.

**Figure 6.**
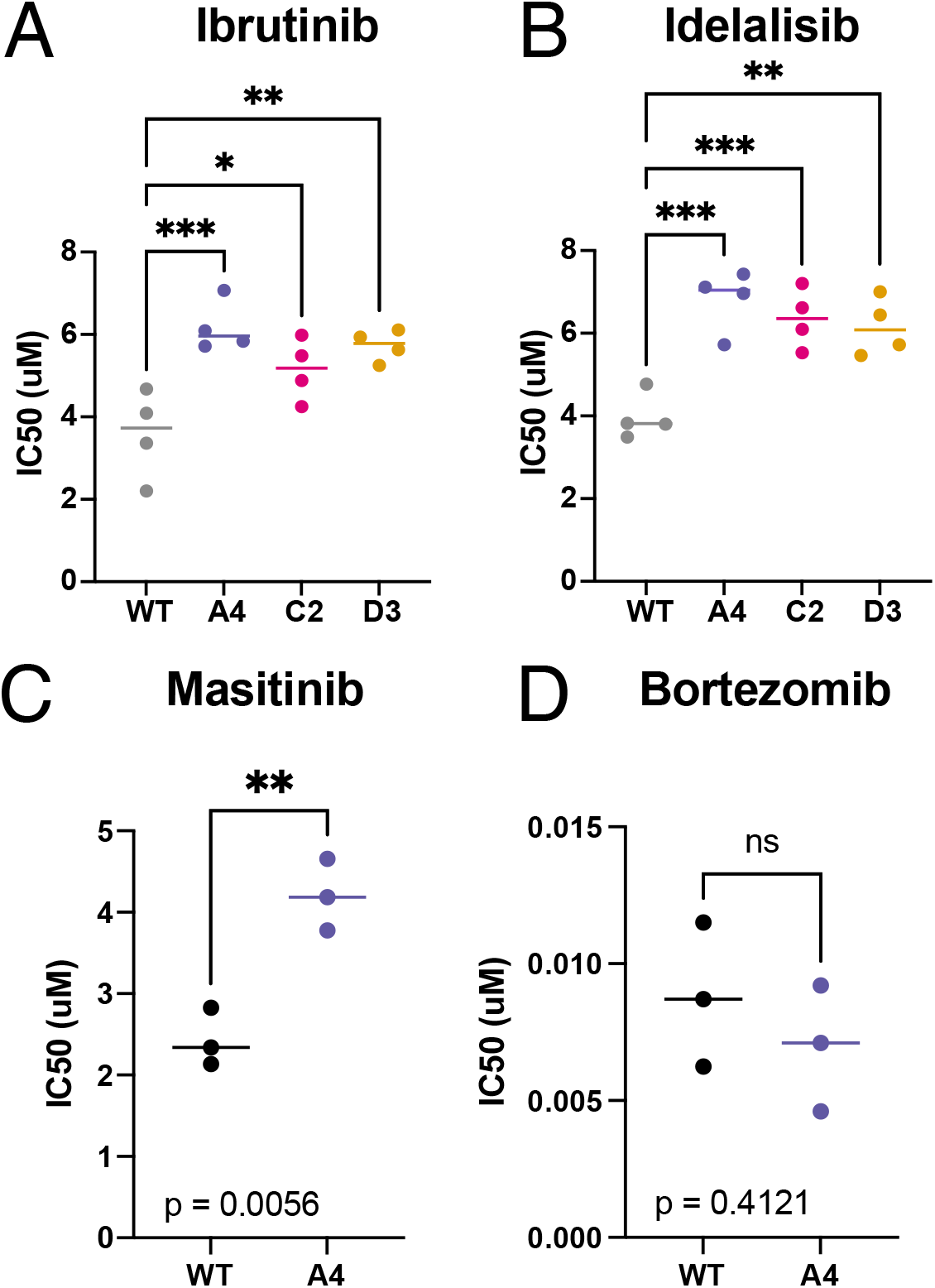
CRISPR-mutant cell lines response to DLBCL targeted therapeutics. Comparison of IC50 for WSU-DLCL2 wild-type and NFKBIZ CRISPR-mutant (A4, C2 and D3) cell lines for (**A**) Ibrutinib, (**B**) Idelalisib, (**C**) Maisitinib, and (**D**) Bortezomib. All points represent the IC50 calculated from independent drug dose-curve assays. All data represent the mean of three or four independent experiments ± SD. Significance was calculated by Student’s t-test or one-way ANOVA (*p ≤ 0.05; **p ≤ 0.01; ***p ≤ 0.001).

**Figure 7.**
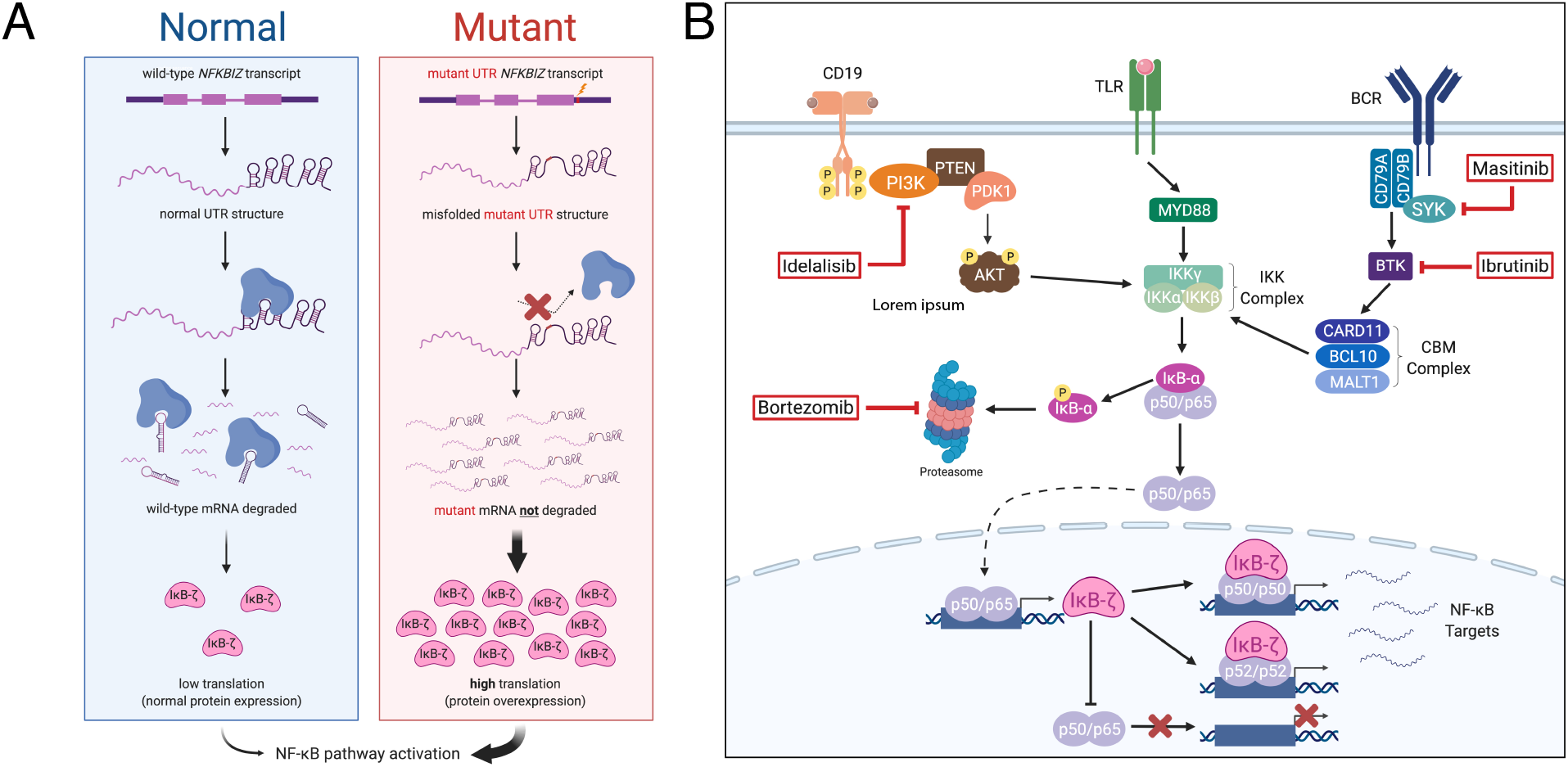
*NFKBIZ* overview models. (**A**) Proposed mechanism of non-coding 3′ UTR mutations in *NFKBIZ* and their effect on mRNA and protein expression and activation of the NF-kB signaling pathway. (**B**) Pathway diagram highlighting where IκB-ζ acts within the NF-κB signaling pathway and the area of the pathway that targeted therapeutics act.

## Discussion

Here, we have demonstrated that *NFKBIZ* 3′ UTR mutations provide a selective advantage to DLBCL and primary B cells *in vitro* and *in vivo*, firmly establishing these non-coding mutations as driver mutations. Through re-analysis of multiple large data sets, we have clarified the prevalence and pattern of *NFKBIZ* 3′ UTR mutations. Although these mutations have not been described in other B-cell malignancies, we observed rare examples in a small minority of FL and BL samples based on whole genome sequencing data, and we note a recent report of a similar pattern of *NFKBIZ* 3′ UTR mutations in T cell lymphoma^53^. This further highlights the importance of elucidating the functional role of IκB-ζ and mutations that affect its expression in hematologic cancers.

Our previous study demonstrated that DLBCL cell lines harbouring *NFKBIZ* 3′ UTR mutations or amplifications generally exhibited higher levels of *NFKBIZ* mRNA and protein^54^. The additional data presented here validates that *NFKBIZ* 3′ UTR mutations cause elevated *NFKBIZ* mRNA and protein expression and provides insights into the mechanism (**Figure 7A**). Isogenic mutant DLBCL cell lines exhibit a consistent selective growth advantage both *in vitro* and *in vivo* and introduction of these mutations into primary GC B-cells provides these cells a growth advantage. This is consistent with the notion that higher IκB-ζ levels resulting from these non-coding mutations promotes cell growth and survival. The most striking effect of these mutations was seen in the WSU-DLCL2 line, which normally exhibits low basal IκB-ζ expression. Despite a more subtle effect in U2932, we confirmed the induction of IκB-ζ expression caused by 3′ UTR mutations. The higher basal expression of IκB-ζ in U2932 WT cells may account for the less dramatic change in expression between mutants and WT compared to WSU-DLCL2 clones. Alternatively, this may be influenced by the nature of the mutations affecting SL3 in U2932, they may exert a different effect than the more common SL2 mutations. Although the RNA change was less obvious in these lines, there was an increase in protein expression, which may indicate that SL3 mutations affect translational efficiency rather than mRNA stability.

The growth advantage afforded to human primary germinal center B-cells by forced overexpression of *NFKBIZ* in *ex vivo* culture confirmed that this gene can confer a growth advantage to non-malignant cells without reliance on other common DLBCL mutations. This result is not always achieved when expressing oncogenes in primary GC B-cells, with many requiring additional perturbations to allow cells to tolerate oncogene expression. Taken together, our data confirms that these non-coding mutations can act as *bona fide* drivers and further suggests that *NFKBIZ* could potentially arise as an early driver of lymphomagenesis.

Currently, LymphGen is the only publicly available approach to classify DLBCLs using genetic features. By comparing the LymphGen assignments of *NFKBIZ*-mutant tumours, we determined that *NFKBIZ* 3′ UTR mutations tend to associate with the BN2 subgroup which corresponds to the other recently described C1^5^ and NOTCH2^7^ DLBCL classes. This separates them from MYD88^L265P^ mutations which are features of the MCD (or C5/MYD88) class, which is consistent with our data showing these mutations are mutually exclusive. These data strongly support the necessity to include *NFKBIZ* mutations as features in future refinements of DLBCL genetic classification systems. We speculate that this would allow appropriate classification of the unclassified cases bearing these mutations, specifically into the BN2/C1/NOTCH2 subgroup. As these classifiers can leave up to 35% of cases unclassified, the inclusion of non-coding mutations could aid in increasing classification rates^8^. As the BN2/C1/NOTCH2 subgroups are known to be enriched for *NOTCH2* and BCR-dependent NF-κB signalling, it can be argued that *NFKBIZ* mutations that deregulate NF-κB targets are a reasonable addition to this group. The BN2/C1/NOTCH2 groups are also known to exhibit genetic alterations affecting immune escape and evasion. The discovery that PD-L1 expression is affected by *NFKBIZ* 3′ UTR mutations suggests *NFKBIZ* could not only be contributing to the NF-κB part of this signature, but also the immune escape. Further exploration of the interplay between these mutations and DLBCL biology will become more feasible only once the mutation status of this region becomes routinely determined through targeted sequencing panels including this region, exome designs with UTR baits, or through whole genome sequencing.

Our analysis identified both PD-L1 and HCK as candidate targets of IκB-ζ, each with potential clinical relevance. Immunotherapies have gained traction in recent years to treat DLBCLs, however their efficacy has been largely dependent on which patients express the proteins relevant for this type of therapy^38,39^. PD-L1 upregulation in response to *NFKBIZ* 3′ UTR mutations could suggest a group of patients who could benefit from this type of treatment, specifically anti-PD1 or PD-L1 antibody therapy^55^. HCK is a src family protein kinase that is expressed in hematopoietic cells^56^. The mutated form of *MYD88* has been shown to trigger transcription and activation of HCK in Waldenström macroglobulinemia^57^, a disease that shares genetic and biological features with ABC DLBCL. HCK has been discussed as a possible therapeutic target for small molecule inhibitors in multiple cancers^58^ and high expression of HCK has been associated with poor prognosis in mantle cell lymphoma^59^. Overexpression of HCK in other cancers can contribute to epithelial mesenchymal transition (EMT), hypoxia and TGFβ signaling^60^. All of these pathways were overexpressed in *NFKBIZ* CRISPR-mutants based on GSEA analysis, consistent with HCK and downstream pathways as relevant targets of IκB-ζ. Knockdown of HCK has been shown to inhibit cell viability, migration and tumor growth^60^. Given the *in vitro* survival benefits afforded by *NFKBIZ* 3′ UTR mutations in cell lines, it is conceivable that HCK expression is partially responsible for these changes and further exploration of its role in DLBCL is warranted.

Induction of IκB-ζ expression through *NFKBIZ* 3′ UTR mutations has a differential effect on the sensitivity of DLBCL cell lines to targeted therapies (**Figure 7B**). The enhanced resistance of these model cell lines to three of the four therapeutics suggests that *NFKBIZ* mutations could be informative as a predictive biomarker for investigational therapies. We were able to identify *NFKBIZ* mutations in patients from a small cohort of nine DLBCL patients treated with ibrutinib monotherapy. Strikingly, all patients harbouring *NFKBIZ* 3′ UTR mutations showed no response to ibrutinib monotherapy^61^. This highlights the pressing need for additional studies to explore the *NFKBIZ* mutation as a component of correlative analyses in clinical trials. This underscores the requirement to not only determine which pathways are mutated in a cancer, but also the location and role of the mutated protein in the affected pathway relative to the protein targeted by individual therapeutics.

## Methods

### Sequencing data re-analysis

Targeted, exome and whole genome sequencing data for the Arthur cohort were combined from previously published papers^9,62,63^. Whole exome sequencing data from Chapuy *et al*.^5^ and Schmitz *et al*.^6^ was obtained from the dbGAP (phs000450) and National Cancer Institute Genomic Data Commons (NCICCR-DLBCL), respectively, and reanalyzed using a standardized variant calling workflow. For Chapuy samples with matched constitutional exomes available, candidate small insertions and deletions were identified using Manta^64^, and these candidate events were provided to Strelka2^65^ to identify both single nucleotide variants and small indels. Passed variant calls were annotated using vcf2maf (https://github.com/mskcc/vcf2maf), and further filtered to remove low quality events using the following filtering criteria: 1) Min read depth >10, 2) Base quality bias p-value >0.01, as determined by comparing bases supporting the reference and alternate alleles using a t-test, 3) Strand bias p-value >0.01, determined by comparing reads supporting the reference and alternate alleles using a fisher’s exact test, 4) Mean mapping quality >50 for reads supporting the alternate allele. 16 samples with excessively high DNA damage were excluded from downstream analysis.

For samples lacking a matched normal, an unpaired constitutional genome was used instead, and candidate small insertions and deletions were identified using Manta^64^ and provided to Strelka2^65^ to identify candidate somatic variants. Unfiltered variant calls (not selecting “PASS” variants) were converted to BED format and provided to MuTect2^66^ to obtain high-quality somatic variant calls, leveraging a panel of normals generated from 58 unrelated constitutional samples. Variant calls were annotated using vcf2maf, and post-filtered as described above. We also filtered any variant with a population allele frequency > 0.005 in any population, as reported in GenomAD^67^, and removed variants with a variant allele frequency < 0.01. Schmitz somatic variant calls were converted to hg19 alignments using CrossMap^68^.

Poly(A) RNA-Seq BAMs from Schmitz *et al*.^6^ was obtained from the National Cancer Institute Genomic Data Commons (NCICCR-DLBCL). These samples were converted back into FASTQs and realigned against the human reference genome GRCh37 using STAR (star ref). Duplicate reads were identified using Picard MarkDuplicates (“Picard Toolkit.” 2019. Broad Institute, GitHub Repository. http://broadinstitute.github.io/picard/; Broad Institute), and gene-level expression data was calculated for each sample using FeatureCounts^69^, using transcript information from Ensembl V87. Allelic imbalance was calculated between DNA and RNA as described previously^54^.

### Cell Culture and Reagents

The cell lines WSU-DLCL2 was purchased from DSMZ and U-2932 was provided by M. Dyer (University of Leicester, UK) and authenticated using short tandem repeat (STR) typing (TCAG, SickKids, Toronto). All DLBCL cell lines were cultured in RPMI 1640 medium (Invitrogen) supplemented with 10% FBS. Protein was extracted from 5⨯10^6^ cells with Pierce RIPA buffer (Thermo, 89901) containing proteasome inhibitor (Sigma, P8340) and quantified using the Pierce BCA Protein Assay Kit (ThermoFisher). 25ug of protein lysate was resolved on a 4-12 % NuPAGE Novex Bis-Tris, 1.0mm Mini Protein Gel (Invitrogen, NP0326BOX) and transferred to a nitrocellulose membrane by wet transfer using the Trans-Blot turbo transfer pack (Bio-Rad). Antibodies for IκB-ζ (Cell Signaling, 9244S), HCK (E1I7F) (Cell Signaling, 14643S), and GAPDH (14C10) (Cell Signaling, 2118S) were diluted according to manufacturer’s recommendations (1:1000). Anti-rabbit IgG HRP conjugate (Promega, W401B) was used to visualize bands with Amersham ECL Western Blotting Detection Reagents (Cytiva, RPN2209) on a Chemidoc digital imager (Bio-Rad).

### CRISPR Gene Editing

Cell lines were transfected using the Amaxa nucleofector with the IDT Alt-R CRISPR/Cas9 system following manufacturer’s recommendations. Briefly, crRNAs were designed using the IDT custom design tool to target the *NFKBIZ* 3′ UTR (SL2: AGCAACACTCACTGTCAGTT and SL3: ATAGACCATTTGCCTTATAT). 2×10^6^ cells per line were electroporated with the generated RNP complex in nucleofector solution V (program X-001). Cells were grown to confluence, then single cell expanded in 96-well plates in 100uL MethoCult H4435 (StemCell). Mutations in clones were verified by Sanger Sequencing (GeneWiz) with primer: CCACATTGGCCATAAGAAAT. Wild-type cells were single cell expanded in the conditions as CRISPR cells to obtain single-cell expanded WT clones.

### Droplet Digital PCR (ddPCR)

RNA was extracted from 5×10^6^ cells per sample using the AllPrep DNA/RNA/miRNA Universal Kit (QIAGEN) according to manufacturer’s recommendations with additional DNase I treatment for RNA extraction. RNA concentration was determined by NanoDrop and 800ug of RNA was used as input for cDNA synthesis using the AB High-Capacity cDNA Synthesis Kit (Applied Biosystems). cDNA was diluted 1:10 for used in ddPCR assays. Custom designed ddPCR assays were used to assess gene expression of NFKBIZ and TBP (control gene) as described before^9^. All plots represent ddPCR assays run on three biological replicates of extracted RNA. Data was analyzed using GraphPad Prism 9 (GraphPad Software).

### T1 RNase Probing Assays

#### *In vitro* transcriptions

In vitro transcriptions were carried out in a mixture of 1 μM dsDNA template, 8 mM GTP, 2 mM UTP, 5 mM CTP and ATP in T7 buffer (2.5 mM Spermidine, 26 mM MgCl2, 10 mM DTT, 0.01% Triton X-100, 40 mM Tris-HCl pH 8.0 at 25 °C) and 1.5 U/μL of T7 RNA polymerase (Applied Biological Materials). Reaction mixtures were incubated at 37°C for 2 h. Transcribed products were made up to 45% formamide, 10 mM EDTA, heated (95°C for 5 min) and subsequently purified by denaturing PAGE, eluted in 0.3 M NaCl and precipitated in 70% ethanol prior to use.

#### CIP 5’ Dephosphorylation of RNA

5’ dephosphorylation of RNA was carried out in a mixture of 1 pmol of RNA 5’ ends in 50 mM Potassium Acetate, 10 mM Magnesium Acetate, 100 µg/ml BSA, 20 mM Tris-acetate (pH 7.9 at 25°C) and 1 U/µl Alkaline Phosphatase, Calf Intestinal (CIP) (New England BioLabs). Reaction mixtures were incubated at 37°C for 30 min. Reactions were stopped by adding equivolume 90% formamide, 20 mM EDTA, heated (95°C for 5 min) and subsequently purified by denaturing PAGE, eluted in 0.3 M NaCl and precipitated in 70% ethanol prior to further use.

#### T4 PNK 5’ Radiolabeling of RNA

Dephosphorylated RNA was 5’ radioactively labeled with T4 Polynucleotide kinase (PNK) in a mixture of 50 pmol of RNA 5’ termini, 20 pmol of [γ-32P] ATP in 10 mM Magnesium Chloride, 5 mM Dithiothreitol (DTT), 70 mM Tris-Hydrochloride (Tris-HCl, pH 7.6 at 25°C) and 1 U/µl T4 PNK (New England BioLabs). Reaction mixtures were incubated at 37°C for 30 min, stopped by heat inactivation, purified by denaturing PAGE and ethanol precipitated as described above, before further use.

#### Alkaline Hydrolysis ladder

5 pmol of 5’ radiolabeled RNA were incubated in 50 mM Sodium bicarbonate (NaHCO_3_) at 90°C for 15 min. Reactions were stopped with 100 mM Tris-HCl and equivolume 90% formamide, 20 mM EDTA, heated (95°C for 5 min) and resolved on 6-10% sequencing denaturing PAGE. Gels were analyzed using a GE Healthcare Amersham Typhoon scanner.

#### Denaturing T1 Digestion

5 pmol of 5’ radiolabeled RNA were incubated in a mixture of 6 M Urea, 20 mM Sodium Citrate (pH 5 at 25°C) and 0.2 U/µl RNase T1 (ThermoFisher). Reaction mixtures were incubated at 50°C for 10 min prior to being stopped by flash freezing in liquid nitrogen for 5 min and adding equivolume 90% formamide, 20 mM EDTA and heated (95°C for 5 min). Samples were resolved on 6-10% sequencing denaturing PAGE and gels were analyzed using a GE Healthcare Amersham Typhoon scanner.

#### Native T1 Digestion

5 pmol of 5’ radiolabeled RNA were incubated in a mixture of 140 mM Potassium Chloride, 1 mM Magnesium Chloride, 0.05% Tween-20, 10 mM Sodium phosphate (pH 7.2 at 25°C) and 0.2 U/µl RNase T1 (ThermoFisher). Reaction mixtures were incubated at 24°C for 10 min or at 37°C for 4 min. Reactions were stopped by flash freezing in liquid nitrogen for 5 min prior to adding equivolume 90% formamide, 20 mM EDTA, heating (95°C for 5 min) and being resolved on 6-10% sequencing denaturing PAGE. Gels were analyzed using a GE Healthcare Amersham Typhoon scanner.

### Competitive Growth Assays

Cell lines were pooled together in equal amounts and grown in culture over 8-10 passages, cells were passaged every 2-3 days when confluence was reached. At each passage a cell pellet was saved and used for DNA extraction with the AllPrep DNA/RNA/miRNA Universal Kit (QIAGEN). 150ng of DNA was used to make libraries with the QIAseq FX DNA Library Kit (QIAGEN). DNA libraries were pooled together for hybridization capture of the *NFKBIZ* gene region using the protocol previously described^9^ based on the XGen hybridization capture workflow from IDT. Probes covering the *NFKBIZ* gene and UTR were used from Arthur *et al*^9^. Sequencing was performed on the Illumina MiSeq, data was analyzed using samtools and bwa-mem. Reads corresponding to each of the WT or mutant clones was counted and used to determine the proportion of each WT/mutant clone at each time point.

### Mouse Xenografts

All cell lines were verified to be mycoplasma negative using the Venor GenM Mycoplasma Detection kit (Sigma Aldrich). 4×10^6^ cells (WSU-DLCL2 WT, A4 or an equally mixed pool of WT/A4/C2/D3 lines) were diluted 1:1 in Matrigel HC and subcutaneously injected into the back of female NSG mice in a volume of 100uL. NSG mice were purchased from an in-house source (Animal Resource Centre, BC Cancer, Vancouver, BC). This study (A14-0259) was approved by the Institutional Animal Care Committee (IACC) at the University of British Columbia/BC Cancer, conducted in accordance with the Canadian Council on Animal Care Guidelines. This study was approved by the Research Ethics Board of the University of British Columbia and the BCCRC, conducted in accordance with the Declaration of Helsinki. Four mice per group were used. Tumor were measured twice weekly. Mice were observed daily and euthanized when they appeared in poor health or when tumors grew to a maximum of 800 mm^3^. Tumor volumes were calculated according to the equation: length × width × height / 2. Xenograft tumors were immediately stored in RNAlater (Sigma-Aldrich) after excision. DNA was extracted following the “Simultaneous purification of Genomic DNA and total RNA, including miRNA, from tissues” protocol for the AllPrep DNA/RNA/miRNA Universal Kit (QIAGEN). The entire tumor was homogenized using the TissueRuptor system from QIAGEN and 600uL of homogenized tissue was used for DNA extraction. DNA library preparation and capture were performed as descried above for competitive growth assays and analyzed similarly.

### RNA-Sequencing and Analysis

RNA was extracted from cell lines as described above for ddPCR assays. Poly(A) RNA libraries were prepared and sequenced by the Genome Sciences Centre in Vancouver. Qualities of total RNA samples were determined using an Agilent Bioanalyzer RNA Nanochip or Caliper RNA assay and arrayed into a 96-well plate (Thermo Fisher Scientific). Polyadenylated (PolyA+) RNA was purified using the NEBNext Poly(A) mRNA Magnetic Isolation Module (E7490L, New England Biolabs) from 500 ng total RNA according to manufacturer’s instructions. First-strand cDNA was synthesized from heat-denatured purified mRNA using a Maxima H Minus First Strand cDNA Synthesis kit (Thermo-Fisher, USA) and random hexamer primers at a concentration of 200 ng/µL along with a final concentration of 40 ng/µL Actinomycin D, followed by PCR Clean DX (Aline Biosciences) bead purification on a Microlab NIMBUS robot (Hamilton Robotics, USA). cDNA was fragmented to by Covaris LE220 sonication to achieve 250-300 bp average fragment lengths. The paired-end sequencing library was prepared following Canada’s Michael Smith Genome Sciences Centre at BC Cancer’s strand-specific, plate-based library construction protocol on a Microlab NIMBUS robot (Hamilton Robotics, USA). DNA quality was assessed with Caliper LabChip GX for DNA samples using the High Sensitivity Assay (PerkinElmer, Inc. USA) and quantified using a Quant-iT dsDNA High Sensitivity Assay Kit on a Qubit fluorometer (Invitrogen) prior to library pooling and size-corrected final molar concentration calculation for Illumina HiSeq sequencing with paired-end 75 base reads. RNA-seq reads were pseudo-aligned using Salmon^70^ (version 0.8.6) to GRCh37. The tximport^71^ Bioconductor R package was then used to summarize transcript level read counts at the gene level. The DESeq2^72^ Bioconductor R package was used to correct the read counts for library size and to obtain differentially expressed genes between conditions of interest employing a threshold of abs(log2FoldChange) > 0.585 and p < 0.05. DESeq2 results were then fed into the FGSEA^73^ Bioconductor R package for pathway enrichment analyses, with a threshold of p < 0.05 set for enriched pathways.

### Drug Treatments

50uL of WSU-DLCL WT or CRISPR-mutant cells were seeded in 96-well plates at a concentration of 5⨯10^5^ cells/ml to be diluted to a final concentration of 2.5⨯10^5^ cells/mL. Cells were then treated with Ibrutinib (PCI-32765 – Selleckchem), Idelalisib (HY-13026/CS-0256 – MedChemExpress), Masitinib (M1838 - AbMole BioScience), or Bortezomib (EMD Millipore Corp) using a ten-point dose titration scheme from 0.01uM to 100 µM, 0.001 µM to 10 µM, 0.01 µM to 100 µM, or 0.001 µM to 10 µM respectively. After 48hr, cell viability was assessed using colorimetric WST-1 reagent (Roche, 11644807001) as per manufacturer’s recommendations. All experimental points were a result of three technical replicates, and all experiments were repeated at least three times. The data was analyzed using GraphPad Prism 9 (GraphPad Software).

Datapoints were normalized to an untreated control and the curves were fitted using a non-linear regression model with a sigmoidal dose response. Each plot shows the IC50 calculated for 3 or 4 biological replicates with mean and standard deviations shown. One-way ANOVA with Dunnet’s multiple comparison’s test was used to compare WT to mutant IC50s for Ibrutinib and Idelalisib. Unpaired 2-tailed student’s t-test was used to compare the IC50s for Masitinib and Bortezomib.

### Plasmid for NFKBIZ Overexpression

The coding sequence of NFKBIZ was amplified by PCR of cDNA from human germinal center B cells and cloned into pMSCV-IRES-GFP II (pMIG II) (addgene #52107) using BamHI and XhoI.

### Primary B-Cell Culture and transduction

Primary GC B cells were purified from fresh tonsil tissue, sourced from Cambridge University Hospitals Trust. Written informed consent of the patient/parent/guardian. Ethical approval for the use of human tissue was granted by the Health Research Authority Cambridgeshire Research Ethics Committee (REC no. 18/EE/0199). The YK6-CD40Lg-IL21 feeder line and protocol for expansion and transduction of human germinal center B cells have been described previously^74,75^. Briefly, germinal center B cells were purified using the human B cell isolation kit (Miltenyi), modified to include negative selection antibodies to IgD and CD34. Cells were then cultured on irradiated YK6-CD40Lg-IL21 feeders and transduced 2-4 days later using GaLV-pseudotyped retrovirus. Retrovirus was produced by transfection of Lenti-X 293T cells with packaging constructs and viral expression vector using TransIT-293 (Mirus). Viral supernatant was harvested 48hr later transfection and filtered through a 0.45 μM filter. Cells were transduced by centrifugation (1500 × *g*, 90 min at 32 °C) with the addition of 10 μg/ml Polybrene and 25 μM HEPES. The viral supernatant was then replaced with fresh RMPI. The frequency of transduced cells was tracked over time by flow cytometry.

## Acknowledgements

This study was supported by a Program Project Grant from the Terry Fox Research Institute (1061) and a Large Scale Applied Research Project from Genome Canada (13124), Genome British Columbia (271LYM), Canadian Institutes of Health Research (CIHR) (GP1-155873) and the British Columbia Cancer Foundation (BCCF). R.D. Morin holds an ASH Foundation Junior Scholar award and is a Michael Smith Foundation for Health Research Scholar. D.H. is supported previously by a Clinician Scientist Fellowship from the MSFHR and currently by a CRUK Senior Research Fellowship (RCCFEL\100072). Research in the Hodson laboratory is supported by the Medical Research Council (MR/M008584/1), the Kay Kendall Leukaemia Fund (KKL1144). The Hodson laboratory receives core funding from Wellcome (203151/Z/16/Z) and MRC to the Wellcome-MRC Cambridge Stem Cell Institute and from the CRUK Cambridge Centre (A25117) and from NIHR Cambridge Biomedical Research Centre (BRC-1215-20014). The datasets have been accessed through the NIH database for Genotypes and Phenotypes (dbGaP) (accession phs001444). A full list of acknowledgements can be found in the supplementary note (Schmitz *et al*., PMID: 29641966). The Genomic Variation in Diffuse Large B Cell Lymphomas study was supported by the Intramural Research Program of the National Cancer Institute, National Institutes of Health, Department of Health and Human Services. The dataset from Chapuy *et al*. was accessed through dbGaP (accession phs000450). This work is conducted as part of the Slim Initiative for Genomic Medicine (SIGMA), a joint U.S.-Mexico project funded by the Carlos Slim Health Institute. Figures included this manuscript were created with BioRender.com. The authors wish to acknowledge Canada’s Michael Smith Genome Sciences Centre, Vancouver, Canada for performing RNA-Sequencing of the CRISPR-mutant cell lines. They also gratefully acknowledge the patient donors of samples used herein.

