## Supplemental Tables and Figures for "Non-coding *NFKBIZ* 3′ UTR mutations promote cell growth and resistance to targeted therapeutics in diffuse large B-cell lymphoma"

### Supplemental Tables/Figures:

**Table S1:** NFKBIZ mutation prevalence in the Arthur/Schmitz/Chapuy cohorts.

| | | | **Subtype** | **Cohort** | | |
|---|---|---|---|---|---|---|
| | | **All** | **ABC** | **DLC** | **Schmitz** | **Chapuy** |
| | **# patients** | 1006 | 511 | 330 | 466 | 210 |
| **NFKBIZ Mutation** | UTR | 101 (10) | 66 (13) | 39 (12) | 49 (11) | 13 (6) |
| | Amplification | 86 (8.5) | 66 (12.9) | 23 (7) | 35 (8) | 28 (13) |
| | TOTAL | 174 (17) | 120 (23.5) | 57 (17) | 79 (17) | 38 (18) |

**Table S2:** *NFKBIZ* 3′ UTR mutations in other WGS cohorts.

| Cohort | # Patients | NFKBIZ UTR mutations (%) | Publication |
|---|---|---|---|
| BL_Adult | 81 | 1 (1.2) | Grande_et_al,2019 |
| BL_Pediatric | 124 | 1 (0.8) | unpublished |
| DLBCL_cell_lines | 15 | 2 (13.3) | Morin_et_al,2013 |
| DLBCL_BC | 117 | 11 (9.4) | Arthur_et_al,2018 |
| DLBCL_ICGC | 87 | 11 (12.6) | Hübschmann_et_al,2021 |
| FL_ICGC | 100 | 4 (4) | Hübschmann_et_al,2021 |
| FL_Kridel | 48 | 1 (2.1) | Kridel_et_al,2016 |

**Table S3:** COO and LymphGen classifications of DLBCLs and FLs from other cohorts.

| | | NFKBIZ UTR mutations (%) | DLBCL (%) | FL (%) |
|---|---|---|---|---|
| **COO** | ABC | 13 (41.9) | 13 (54.2) | 0 (0) |
| | GCB | 4 (12.9) | 4 (16.7) | 0 (0) |
| | UNCLASS | 2 (6.5) | 2 (8.3) | 0 (0) |
| | NA | 12 (38.7) | 5 (20.8) | 5 (100) |
| | TOTAL | 31 | 24 | 5 |
| **LymphGen** | BN2 | 13 (41.9) | 9 (37.5) | 3 (60) |
| | EZB | 4 (12.9) | 2 (8.3) | 2 (40) |
| | ST2 | 1 (3.2) | 1 (4.2) | 0 (0) |
| | Other | 9 (29) | 8 (33.3) | 0 (0) |
| | Composite | 2 (6.5) | 2 (8.3) | 0 (0) |
| | NA | 2 (6.5) | 2 (8.3) | 0 (0) |
| | TOTAL | 31 | 24 | 5 |

**Table S4:** LymphGen classifications of all patients and those with *NFKBIZ* mutations in Arthur/Schmitz/Chapuy cohorts.

| | Patients with NFKBIZ Mutation (% Patients) | | | All Patients (% Patients) |
|---|---|---|---|---|
| **LymphGen Classification** | **All** | **UTR** | **Amplifications** | |
| A53 | 35 (19.7) | 10 (9.9) | 28 (32.2) | 103 (10.2) |
| BN2 | 33 (18.5) | 31 (30.7) | 4 (4.6) | 118 (11.7) |
| EZB | 7 (3.9) | 4 (4.0) | 3 (3.4) | 166 (16.5) |
| MCD | 25 (14) | 10 (9.9) | 14 (16.1) | 106 (10.5) |
| ST2 | 1 (0.6) | 1 (1) | 0 (0) | 57 (5.7) |
| N1 | 0 (0) | 0 (0) | 0 (0) | 18 (1.8) |
| Unclass | 65 (36.5) | 37 (36.6) | 32 (36.8) | 357 (35.5) |
| Composite | 12 (6.7) | 8 (7.9) | 6 (6.9) | 81 (8.1) |
| A53/MCD | 3 | 0 | 3 | |
| A53/ST2 | 1 | 1 | 1 | |
| A53/BN2 | 1 | 1 | 0 | |
| A53/EZB | 1 | 1 | 1 | |
| BN2/MCD | 3 | 3 | 0 | |
| BN2/ST2 | 1 | 0 | 1 | |
| EZB/ST2 | 2 | 2 | 0 | |
| **TOTAL** | 178 | 101 | 87 | 1006 |

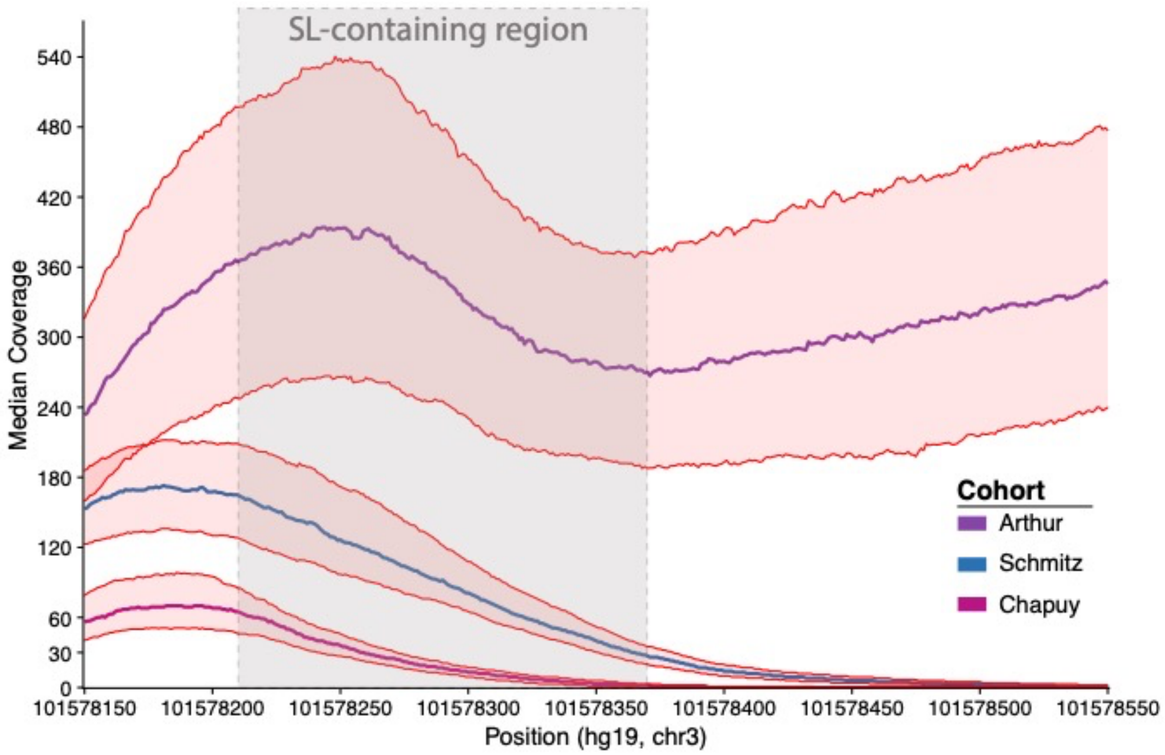


**Figure S1. Coverage of *NFKBIZ* 3′ UTR within each cohort**. Median coverage at each base position across the NFKBIZ 3′ UTR position with the 25th and 75th quartiles shown shaded in red for each of the three cohorts. The region specifically containing the stem-loops structures where most mutations occur is shown in between two vertical dashed lines. The cut-off of 30X coverage is shown by the vertical dashed line, the threshold to be able to confidently call mutations in a region. The Arthur cohort is targeted sequencing with probes for the UTR region. The Schmitz cohort is exome data with mostly sufficient coverage to call mutations, especially early in the UTR region. The Chapuy cohort coverage was very low and not "callable" in many cases, especially in the locations after SL2 (~chr3:101578280).

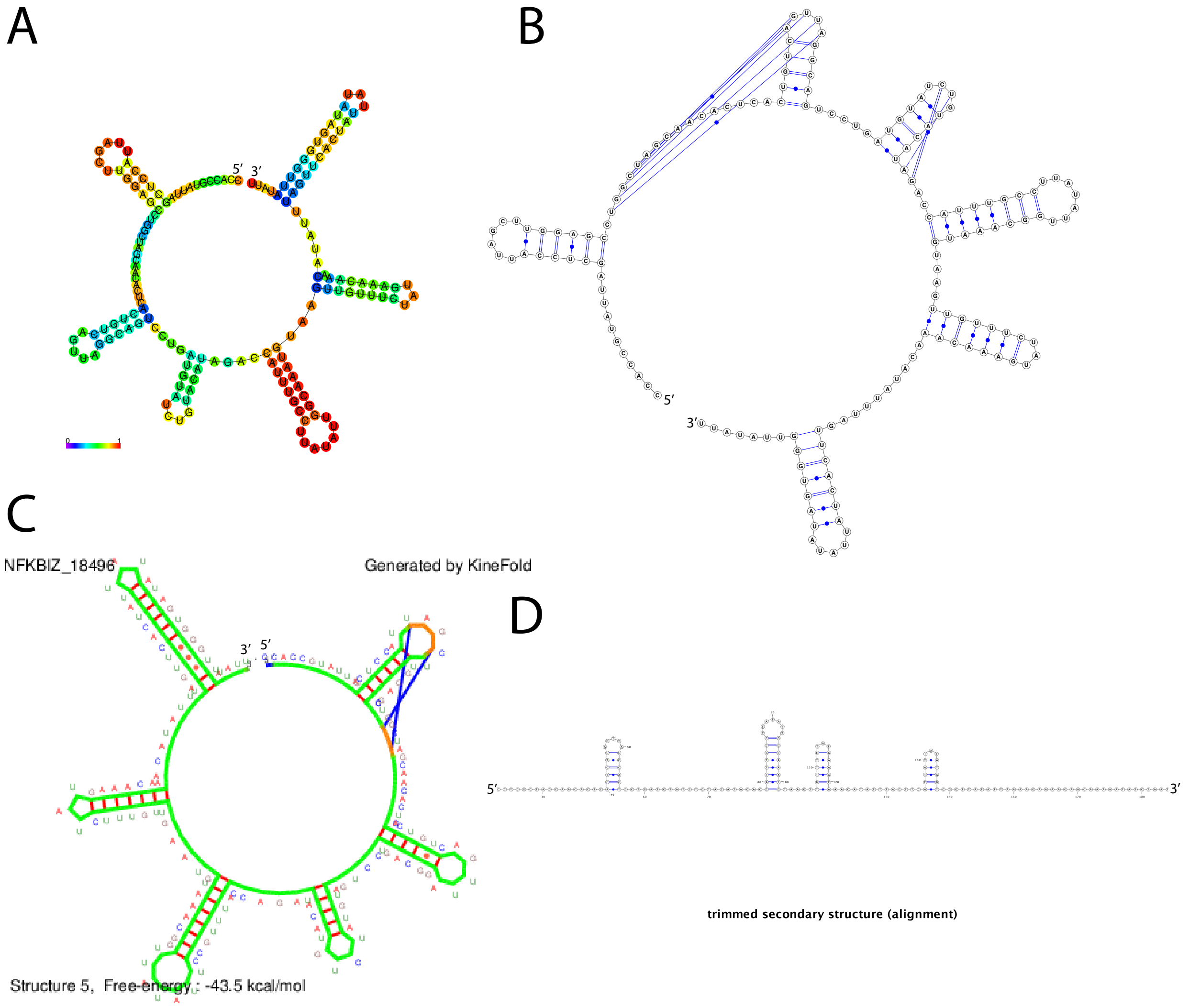


**Figure S2. *In Silico* RNA folding algorithm predictions.** (A) CentroidFold (B) IPknot (C) KineFold (D) RNAalifold. Bases are represented in circles and/or coloured bases, base pairs are represented by lines connecting the bases. Long blue lines in B and C represent possible pseudoknots.

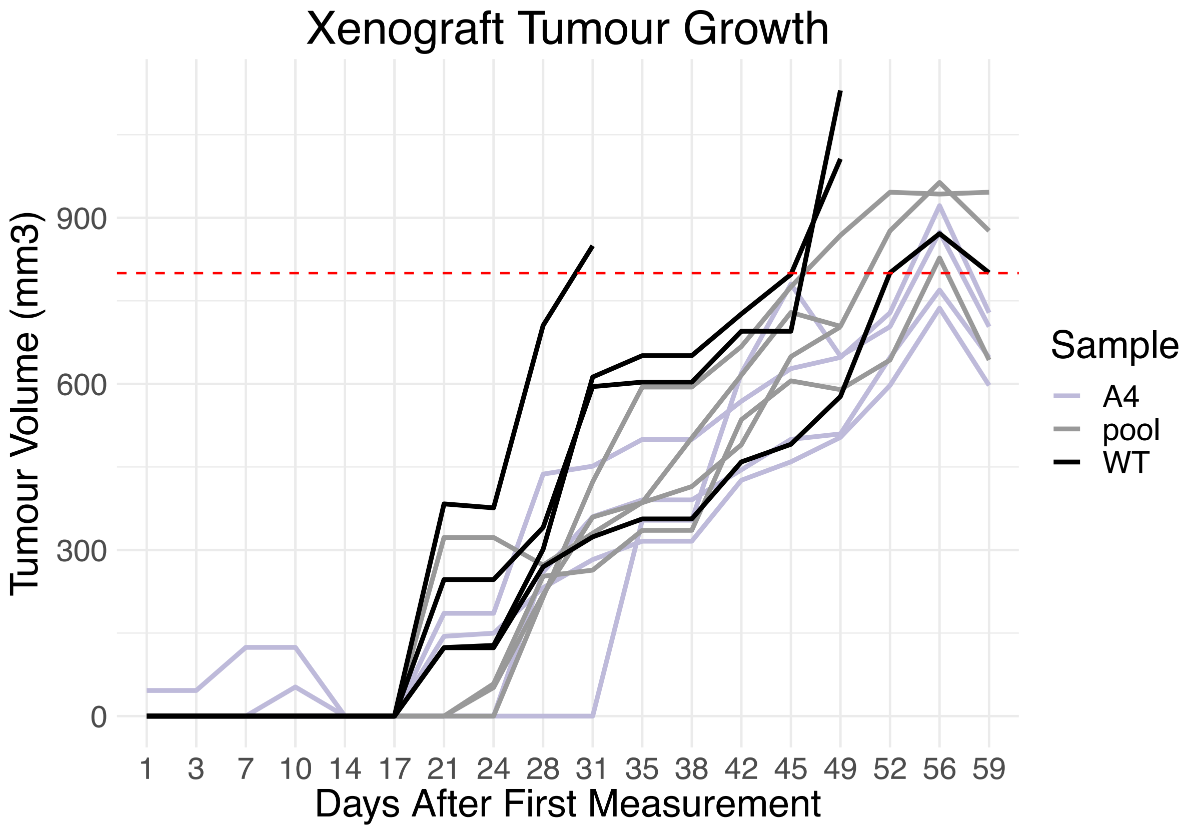


**Figure S3. Growth of xenografted tumors.** Growth rate of tumors from three groups: WT (wild-type WSU-DLCL2 cells, A4: WSU-DLCL2 A4 CRISPR-mutant cells, and pool: pool of WT, A4, C2 and D3 mutant cells). All appeared to grow at a similar rate. End point of experiment is shown by red dashed line ($800mm^3$ tumor size).

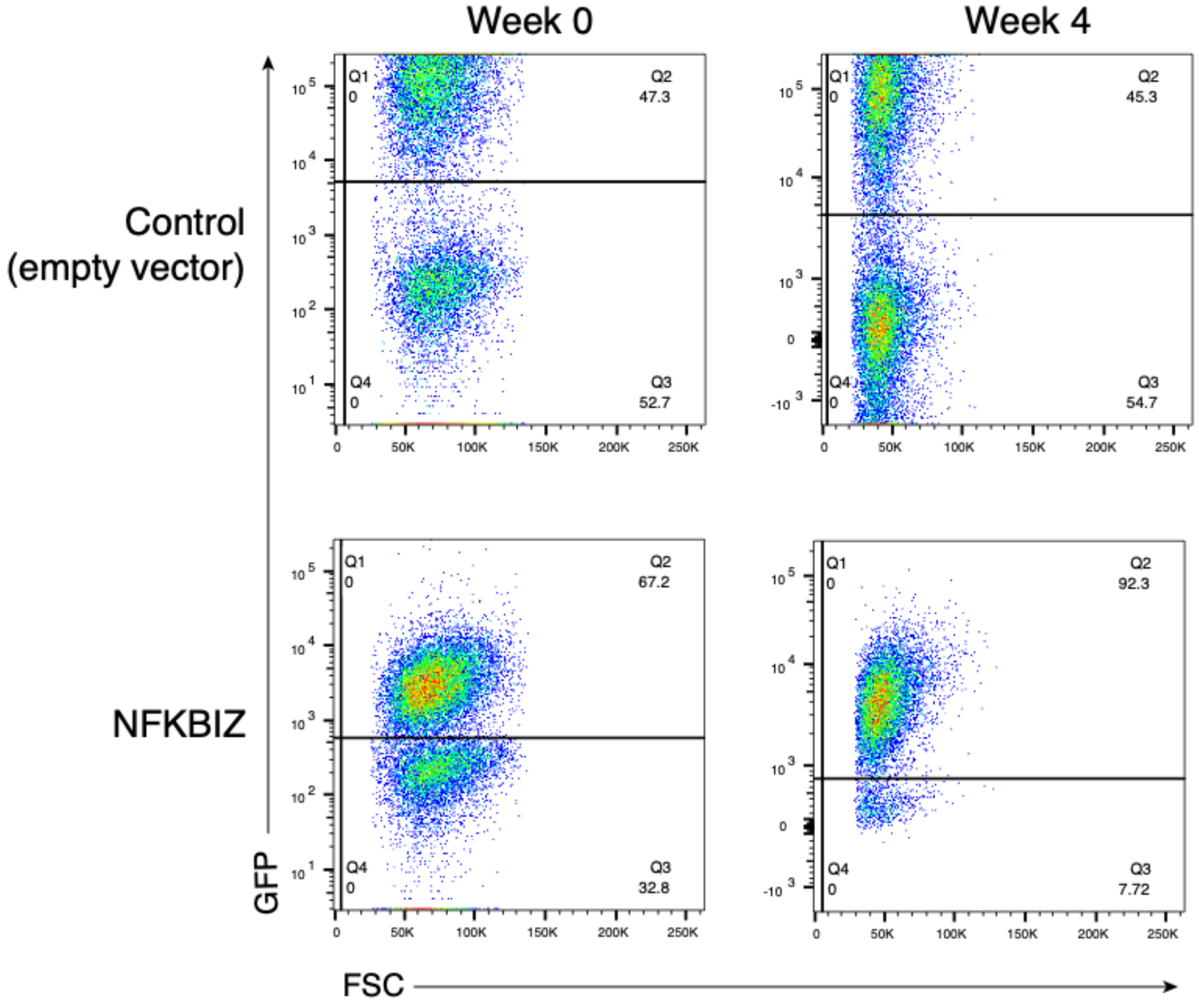


**Figure S4. Forced expression of *NFKBIZ* provides a competitive advantage to human primary germinal center B cells cultured *ex vivo*.** Example flow cytometry plots showing progressive expansion of the GFP-positive population in *NFKBIZ*-transduced but not control cells by comparing percent of GFP cells at week 0 and week 4.

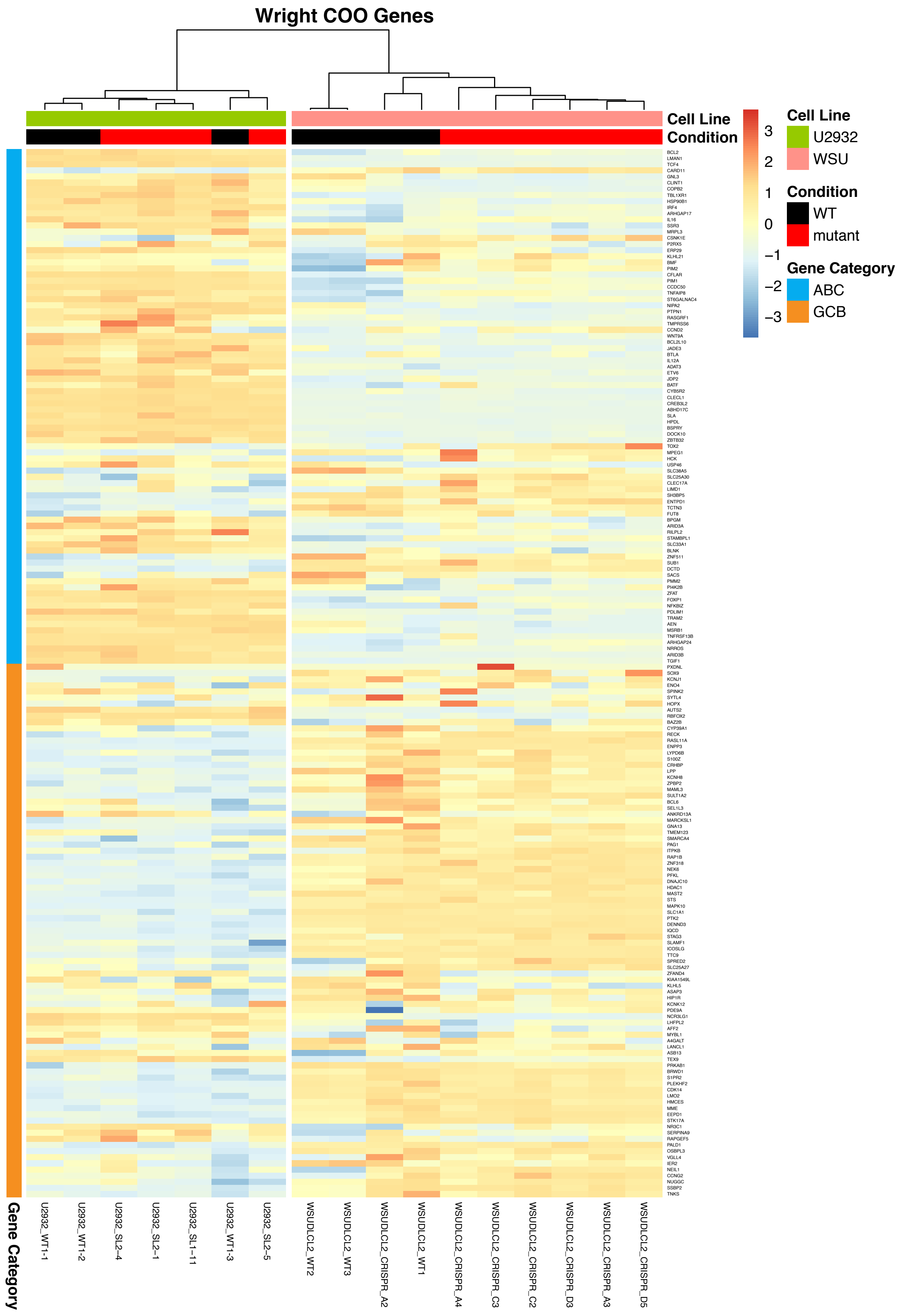

Wright COO Genes
Cell Line
Condition
Cell Line
U2932
WSU
Condition
WT
mutant
Gene Category
ABC
GCB
3
2
1
0
−1
−2
−3
BCL2
LMAN1
TCF4
CARD11
GNL3
CLINT1
COPB2
TBL1XR1
HSP90B1
IRF4
ARHGAP17
IL16
SSR3
MRPL3
CSNK1E
P2RX5
ERP29
KLHL21
BMF
PIM2
CFLAR
PIM1
CCDC50
TNFAIP8
ST6GALNAC4
NIPA2
PTPN1
RASGRF1
TMPRSS6
CCND2
WNT9A
BCL2L10
JADE3
BTLA
IL12A
ADAT3
ETV6
JDP2
BATF
CYB5R2
CLECL1
CREB3L2
ABHD17C
SLA
HPDL
BSPRY
DOCK10
ZBTB32
TOX2
MPEG1
HCK
USP46
SLC38A5
SLC25A30
CLEC17A
LIMD1
SH3BP5
ENTPD1
TCTN3
FUT8
BPGM
ARID3A
RILPL2
STAMBPL1
SLC33A1
BLNK
ZNF511
SUB1
DCTD
SACS
PMM2
PI4K2B
ZFAT
FOXP1
NFKBIZ
PDLIM1
TRAM2
AEN
MSRB1
TNFRSF13B
ARHGAP24
NRROS
ARID3B
TGIF1
PXDNL
SOX9
KCNJ1
ENO4
SPINK2
SYTL4
HOPX
AUTS2
RBFOX2
BAZ2B
CYP39A1
RECK
RASL11A
ENPP3
LYPD6B
S100Z
CRHBP
LPP
KCNH8
ZPBP2
MAML3
SULT1A2
BCL6
SEL1L3
ANKRD13A
MARCKSL1
GNA13
TMEM123
SMARCA4
PAG1
ITPKB
RAP1B
ZNF318
NEK6
PFKL
DNAJC10
HDAC1
MAST2
STS
MAPK10
SLC1A1
PTK2
DENND3
IQCD
STAG3
SLAMF1
ICOSLG
TTC9
SPRED2
SLC25A27
ZFAND4
KIAA1549L
KLHL5
ASAP3
HIP1R
KCNK12
PDE9A
NCR3LG1
LHFPL2
AFF2
MYBL1
A4GALT
LANCL1
ASB13
TEX9
PRKAB1
BRWD1
S1PR2
PLEKHF2
CDK14
LMO2
HMCES
MME
EEPD1
STK17A
NR3C1
SERPINA9
RAPGEF5
PALD1
OSBPL3
VGLL4
IER2
NEIL1
CCNG2
NUGGC
SSBP2
TNKS
Gene Category
U2932_WT1-1
U2932_WT1-2
U2932_SL2-4
U2932_SL2-1
U2932_SL1-11
U2932_WT1-3
U2932_SL2-5
WSUDLCL2_WT2
WSUDLCL2_WT3
WSUDLCL2_CRISPR_A2
WSUDLCL2_WT1
WSUDLCL2_CRISPR_A4
WSUDLCL2_CRISPR_C3
WSUDLCL2_CRISPR_C2
WSUDLCL2_CRISPR_D3
WSUDLCL2_CRISPR_A3
WSUDLCL2_CRISPR_D5

**Figure S5. Expression of genes associated with COO in WSU-DLCL2 and U2932 WT and CRISPR Clones.** Expression of genes used to assess COO status from gene expression profiling is shown for clones derived from the GCB-cell line WSU-DLCL2 and the ABC cell line U2932. ABC associated genes are highly expressed in the U2932 clones and some are highly expressed in the WSU-DLCL2 *NFKBIZ*-mutant CRISPR clones.

### Supplemental Methods:

#### In silico RNA folding algorithms

The same sequence of RNA was used for all prediction methods which contains the end of the NFKBIZ coding sequence and the start of the 3′ UTR where mutations occur most often.

CCACCGUAUUAGCUCCAUUAGCUUGGAGCCUGGCUAGCAACACUCACUGUCAGU
UAGGCAGUCCUGAUGUAUCUGUACAUAGACCAUUUGCCUUAUAUUGGCAAAUGU
AAGUUGUUUCUAUGAAACAAACAUAUUUAGUUCACUAUUAUAUAGUGGGUUAUA
UU

**IPknot.** The web-based application (Version 1.3.1) was used for structural prediction (http://rtips.dna.bio.keio.ac.jp/ipknot/). The above sequence of RNA was used as input with the parameters: Sequence length (164nt), level (2), scoring mode (McCaskill model), Without refining parameters, weight for true base pairs (level1:2; level2:16)

**CentroidFold.** The web-based application was used for structural prediction (http://rtools.cbrc.jp/centroidfold/). The above sequence of RNA was used as input with the parameters: Interface engine (McCaskill(BL)), Weight of base pairs (2^2).

**KineFold.** The web-based application was used for structural prediction (http://kinefold.curie.fr/cgi-bin/form.pl). The above sequence of RNA was used as input with the parameters: Type of Stochastic Simulation (co-transcriptional fold), simulated molecular time (suggested), pseudoknots (allowed), entanglements (non crossing), random seed (18496).

**RNAalifold.** Jalview software was used to analyze multiple sequence alignment (MSA) downloaded from Galaxy for the region specified above. RNAalifold was used to predict the secondary structure from the MSA and the structure for hg19 sequence was performed with VARNA.
